# Friendship modulates physical effort and neural dynamics during group coordination in shared social networks

**DOI:** 10.64898/2025.12.30.695445

**Authors:** Qianliang Li, Aliaksandr Dabranau, Kyveli Kompatsiari, Ivana Konvalinka

## Abstract

Human social interactions are embedded within social networks, yet how existing interpersonal relationships influence the behavioural and neural dynamics of collective action remains unclear. We combined social network mapping with three-person EEG-hyperscanning to investigate how friendship asymmetry within shared social networks modulates physical effort allocation during real-time coordination. Triads consisting of two close friends and one non-friend performed a novel joint force task. Non-friends consistently produced greater and more vigorous force relative to friends, particularly those embedded in smaller classroom social networks. At the neural level, non-friends also showed stronger alpha- and beta-band event-related desynchronization, inter-subject correlation, and group phase synchrony, indicating greater engagement of action-related and attentional processes. These findings suggest that non-friends invest more physical effort and attention when interacting in socially asymmetric group settings, demonstrating how real-world relational context within social networks is dynamically embodied in the neural and behavioural mechanisms supporting human group coordination.

## 1 Introduction

Friendship is one of the most powerful social ties humans form, shaping well-being, identity, and opportunities across the lifespan (Hartup and Stevens, 1997; van der Horst and Coffé, 2012; Dunbar, 2018; Alsarrani et al., 2022; Pezirkianidis et al., 2023; Krems and Kaufman, 2026). It forms the foundation of our social networks, which define the broader societal context within which individuals coordinate everyday behaviour - from moving furniture and cooking together, to group decision making at work and team sports. Such coordinated actions rely not only on cognitive mechanisms, such as integration of sensory information from one’s own and others’ actions, anticipation and adaptation to others’ behaviour, and real-time allocation of effort (Sebanz et al., 2006; Knoblich et al., 2011; Vesper et al., 2017; Sebanz and Knoblich, 2021), but also on the relational context in which coordination unfolds (Tomasello, 2014; Clark-Polner and Clark, 2014; Chierchia et al., 2020). Yet despite this prominent role, we know remarkably little about how friendship dynamics in real-world social groups influence moment-to-moment processes through which people coordinate their actions together and allocate effort to joint goals.

While group interactions create opportunities for unequal contributions to collective tasks, previous research shows that the amount of effort individuals contribute depends on the social conditions under which they interact (Karau and Williams, 1993). Research on social loafing has shown that individuals often reduce their effort when working toward a collective rather than individual outcome, but that this reduction depends on factors such as evaluation of individual contributions and the cohesiveness of the group (Williams and Karau, 1991; Karau and Williams, 1993, 1997; Karau and Wilhau, 2020). Conversely, research on social facilitation shows that the presence of others can also enhance effort and task performance (Zajonc, 1965; Bond and Titus, 1983), particularly when individuals are subject to evaluation by others (Leary and Kowalski, 1990; Izuma, 2012). Effort during collective action is thus sensitive to the social context in which the actions take place. One important aspect of this context is the existing relationship between interacting individuals, such as whether they are friends or not. Greater group cohesiveness and commitment among friends could reduce social loafing and promote greater contributions to a shared outcome (Karau and Williams, 1993, 1997; Wieselquist et al., 1999). At the same time, interactions with less closely affiliated others may carry greater potential for social evaluation and could therefore increase engagement and effort during collective action (Leary and Kowalski, 1990; Izuma, 2012). Taken together, these perspectives yield different predictions about whether friends or non-friends would contribute more effort when acting together toward a shared goal.

Understanding how friendship influences effort allocation during collective action requires paradigms that capture realtime interactions among people with existing relationships, while measuring the behavioural and neural mechanisms through which relational context is expressed during coordination (Hein et al., 2024). Recent advances in social neuroscience and joint action have highlighted the need for interactive multi-person neuroimaging paradigms to uncover how social factors influence interactive behaviour and underlying neural mechanisms (De Jaegher et al., 2010; Dumas, 2011; Konvalinka and Roepstorff, 2012; Schilbach et al., 2013; Dingemanse et al., 2023). Joint action studies show that successful coordination requires temporal prediction of others’ actions and mutual adaptation (Knoblich and Jordan, 2003; Keller et al., 2007; Konvalinka et al., 2010; Noy et al., 2011; Vesper et al., 2011; Pecenka and Keller, 2011; Keller et al., 2014; Vesper et al., 2017; Pezzulo et al., 2019; Sebanz and Knoblich, 2021; Konvalinka et al., 2023). Whether friendship influences these coordination processes, and whether its effects depend on prediction or continuous monitoring of others’ actions, remains largely unknown.

The neural mechanisms supporting these processes are more poorly understood. At the individual-level, enhanced mu-, alpha- and beta-band power suppression and event-related desynchronization (ERD) have been associated with motor prediction, action execution, and increased attentional allocation to others’ behaviour (Tognoli et al., 2007; Perry et al., 2011; Dumas et al., 2012; Naeem et al., 2012; Konvalinka et al., 2014; Zamm et al., 2021; Zimmermann et al., 2022; Flösch et al., 2023). At the inter-brain level, complementary measures have been used to characterize different forms of neural alignment. Inter-brain phase synchronization quantifies phase coupling between two individuals’ neural oscillations and has been observed during action coordination (Dumas et al., 2010; Müller et al., 2013; Gugnowska et al., 2022). Inter-subject correlation (ISC), in turn, quantifies similarity in the temporal evolution of neural responses across individuals (Schilbach and Redcay, 2025). When people attend to the same visual or auditory stimulus, even in the absence of interaction, their neural signals become correlated through common stimulus locking (Dmochowski et al., 2012; Golland et al., 2015; Cohen and Parra, 2016; Nastase et al., 2019; Madsen and Parra, 2022). Higher ISC has thus been purposed to serve as a marker of joint attentional engagement, reflecting more consistent, time-locked processing of the task across individuals (Hasson et al., 2004; Dmochowski et al., 2012, 2014; Cohen et al., 2018; Nastase et al., 2019; Rai et al., 2025; He et al., 2026).

Friendship has also been associated with greater inter-subject neural similarity, both at rest (Hyon et al., 2020b) and during passive video watching (Parkinson et al., 2018; Hyon et al., 2020a; Ma and Liu, 2024; Shen et al., 2025). During passive viewing, greater stimulus-locked neural similarity has been linked to future friendship formation (Shen et al., 2025), and shown to decrease with increasing social distance in real-world social networks (Parkinson et al., 2018; Ma and Liu, 2024; Shen et al., 2025). This neural similarity may reflect more shared ways of perceiving external stimuli and responding to the world, which may in turn facilitate friendship formation (Parkinson et al., 2018; Shen et al., 2025).

ERD and ISC thus provide complementary indices of the neural dynamics that support joint coordination. ERD provides an index for individual sensorimotor prediction and action execution processes, while ISC measures reflect shared stimulus-locked task processing and task engagement. However, whether friendship influences these neural dynamics during active interaction remains unclear (Ma and Liu, 2024) and may differ depending on the task and social context (Ma and Liu, 2024; Speer et al., 2024; Zhou et al., 2025).

Here we investigated how friendship asymmetry within real-world social groups influences real-time collective coordination dynamics, by combining social network mapping with three-person EEG-hyperscanning during interaction. We focused on how friendship dynamics within shared social networks influence physical effort allocation and neural measures of engagement during group coordination. Shared social networks were specifically targeted to study coordination among individuals embedded within a common social environment rather than interactions between strangers. Such networks provide a social context in which individuals may possess implicit knowledge not only of their relationships with others, but also of others’ positions within the larger group, consistent with evidence that people spontaneously encode the social network positions of their classmates (Parkinson et al., 2017).

We mapped social networks across seven bachelor program study lines (Figure 1A), and recruited triads composed of two mutually close friends and one non-friend from the same study line networks. Participants performed a novel three-person coordination task (Figure 1B) - the “Force Game” - in which they jointly coordinated their force contributions to reach specific target forces by applying pressure to individual pressure measuring devices, while receiving continuous visual feedback (wFb) or non-continuous visual feedback (nFb) of joint, but not individual, contributions (Figure 1C). Continuous feedback allowed participants to monitor the evolving joint force and adapt their own contribution throughout the interaction, whereas in the absence of continuous feedback they had to rely on their prediction of the joint force and relative contributions without the possibility of online correction. This manipulation therefore allowed us to investigate whether friendship-related differences in behaviour and neural dynamics depended on adaptation or prediction during coordination. This design allowed us to test the two competing hypotheses: whether friends or non-friends contribute more physical effort during coordination (with and without continuous adaptation), and whether differences in effort are accompanied by differences in task-related attention and engagement.

**Fig. 1.**
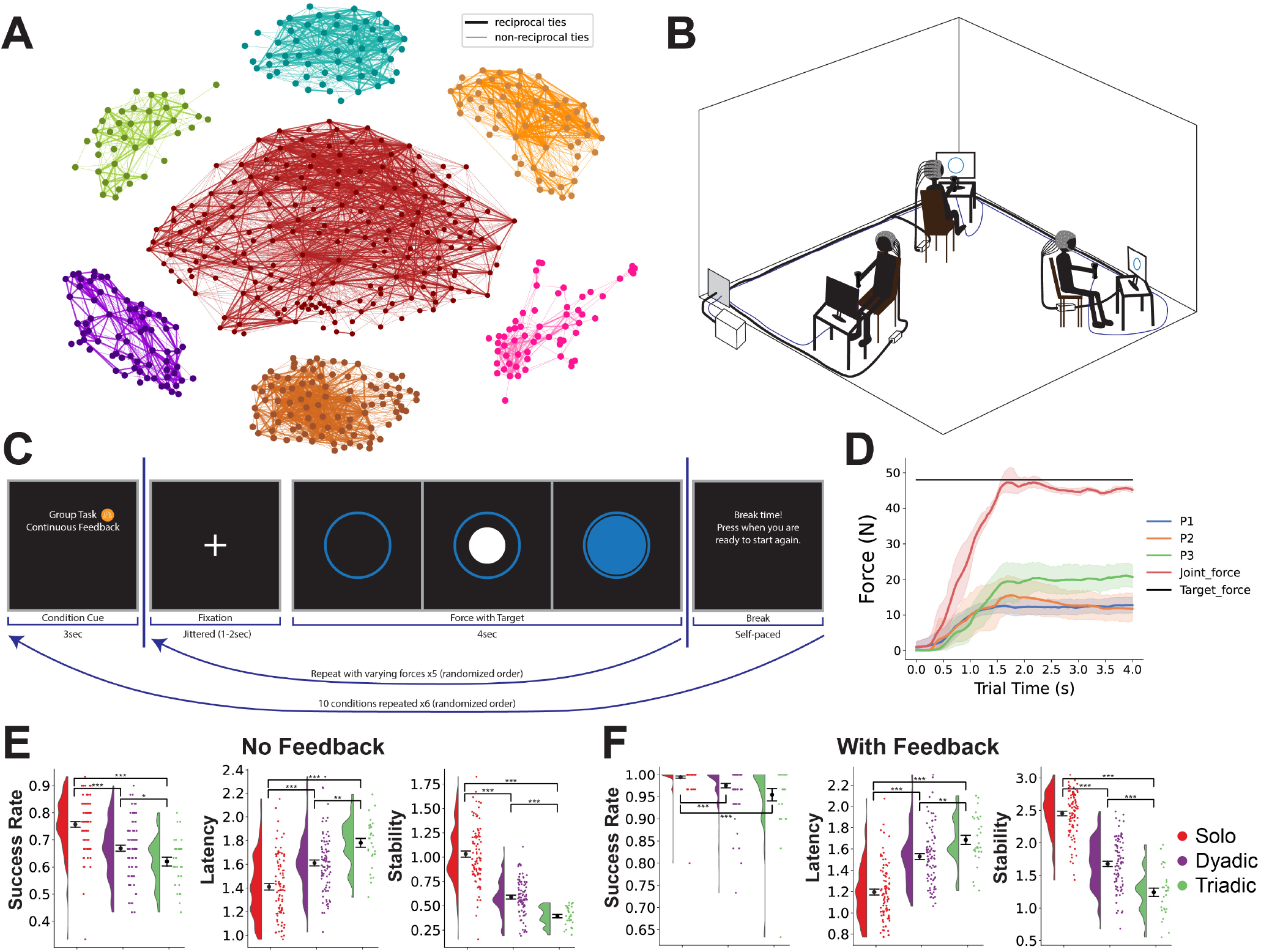
Schematic of the Force Game setup. A) Social networks of the seven mapped bachelor program study lines. B) Triads, consisting of two close friends and one non-friend from the same study line, performed the Force Game together. C) Timeline of the experimental paradigm for a single trial. The blue circle indicates the target force, and the filled white circle indicates the joint force produced, in a trial with continuous visual feedback. The arrows beneath the timeline indicate the repeated loops corresponding to different force levels and conditions. D) Example of the averaged forces across the 4-second triadic trials with continuous visual feedback from one triad. E-F) Behavioural performance, estimated with success rate, latency and stability around the target force, for trials E) without continuous visual feedback and F) with continuous feedback. ^∗^*p <* 0.05, ^∗∗^*p <* 0.01, ^∗∗∗^*p <* 0.001.

Importantly, most prior interactive hyperscanning studies have focused on dyads (Ma and Liu, 2024), whereas real-world coordination often occurs in small groups with existing group relational asymmetries. Studying triads allowed us to introduce a minimal group structure in which asymmetric social relationships can be embedded. Combined with recruiting participants from shared social networks, this design therefore allows us to investigate how naturally occurring friendship asymmetries and relative social position within an existing social structure influences real-time group coordination.

In this study, we found that non-friends contributed greater and more vigorous force than friends during joint tasks. This effect was strongest for non-friends from smaller class social networks, showing that the behavioural effect of friendship asymmetry varied with the broader social environment in which participants were embedded. Non-friends also exhibited greater alpha and beta event-related desynchronization, higher neural inter-subject correlation, and increased neural group phase synchrony (Richardson et al., 2012) -which we used as a phase-based measure of task-aligned neural processing - suggesting that non-friends displayed increased attention and engagement during the task. By combining relative friendship information from shared social networks, real-time behavioural coordination, and multi-brain imaging, we show that asymmetric social relationships are embodied in the motor and cognitive processes supporting real-time human group coordination.

## 2 Results

### 2.1 Social network mapping

We first mapped the social networks of seven bachelor program study lines, with class sizes ranging from 43 to 184 students (*M* = 83.14; *SD* = 44.26). Students were asked to complete a questionnaire in which they checked off the individuals they considered to be friends or close friends (see Supplementary Methods for definitions) from a roster of all of their classmates in the same study line. 378 students gave informed consent to have their social networks mapped across the seven social networks (Figure 1A), ranging from 28 to 113 (*M* = 54.0; *SD* = 27.09) students per study line class.

A subset of the students (*N* = 93) were recruited as triads, consisting of two mutually reported close friends and one non-friend (defined as mutual lack of social ties to the friend-pair), to participate in the Force Game experiment (Figure 1B). These participants filled out an additional friendship questionnaire, where they were asked to list their friends and close friends outside of the study line class social networks, to obtain a more complete estimate of the number of friends for each participant. Overall, the non-friends had significantly fewer friends (Supplementary Figure S1; Class friends: mean difference Δ*M* = 4.65, Cohen’s *d* = 0.511, *p* = 0.032; Class close friends: Δ*M* = 3.08, *d* = 0.781, *p* = 0.002; Friends outside classroom Δ*M* = 2.70, *d* = 0.471, *p* = 0.043; Close friends outside classroom Δ*M* = 3.59, *d* = 0.624, *p* = 0.011; all p-values were adjusted for multiple comparisons using the false-discovery rate (FDR) procedure).

### 2.2 Non-friends contribute greater and more vigorous forces compared to friends in group coordination

Participants were instructed to apply forces by pressing on individual pressure measuring devices in order to individually or jointly reach specific target forces visually shown on their individual screens at the start of each trial (Figure 1C), whilst three-person hyperscanning EEG was measured. Participants either performed the task alone (solo) or interactively. In interactive trials, they had to reach the target force together in triads (referred to as triadic trials), or in dyads where one person’s device did not contribute to the total force (dyadic trials), a manipulation they were not aware of. Additionally, we manipulated whether the interaction recruited adaptive or predictive mechanisms, by either giving continuous visual feed-back - displayed as a filled circle on the screen - or no feedback across the 4-second trial followed by discrete feedback of the reached force at the end of the trial. The feedback given in interactive trials was always of the joint force, hence individual forces had to be inferred. The participants were thus not given feedback of their relative contributions. Each condition was repeated 30 times with varying target forces, represented by the radius of the target circle. An example of the trial averaged produced forces across the 4-second interaction from one triad can be seen in Figure 1D, depicting group interaction with continuous feedback.

To validate the Force Game paradigm, we estimated how well the participants performed the task. As expected, participants performed better with continuous visual feedback and performance deteriorated as more people contributed, i.e., with the best performance observed in solo trials, and worst in triadic ones (Figure 1E-F), likely reflecting greater perceptual demands placed on each participant when integrating the collective joint force.

To test whether different coordination mechanisms emerged between friends and non-friends, we first investigated how much relative force each participant contributed to the joint task using a linear mixed-effects model. Here, a systematic asymmetry emerged: non-friends contributed greater physical force than friends (Figure 2), and the effect of friendship was modulated by group size and feedback, as indicated by a three-way interaction (*β* = 0.028, 95% CI [0.000, 0.056], SE = 0.014, *z* = 1.984, *p* = 0.047). To interpret this interaction, we also examined simple pairwise contrasts between friends and non-friends across conditions, finding that the non-friends consistently contributed more force than friends across all interactive conditions (Dyadic wFb: *β* = 0.066, 95% CI [0.006, 0.125], SE = 0.030, *z* = 2.17, *p* = 0.040; Triadic wFb: *β* = 0.061, 95% CI [0.003, 0.120], SE = 0.030, *z* = 2.05, *p* = 0.040; Dyadic nFb: *β* = 0.109, 95% CI [0.051, 0.168], SE = 0.030, *z* = 3.65, *p* = 0.001; Triadic nFb: *β* = 0.077, 95% CI [0.019, 0.135], SE = 0.030, *z* = 2.59, *p* = 0.019; FDR corrected across all conditions).

**Fig. 2.**
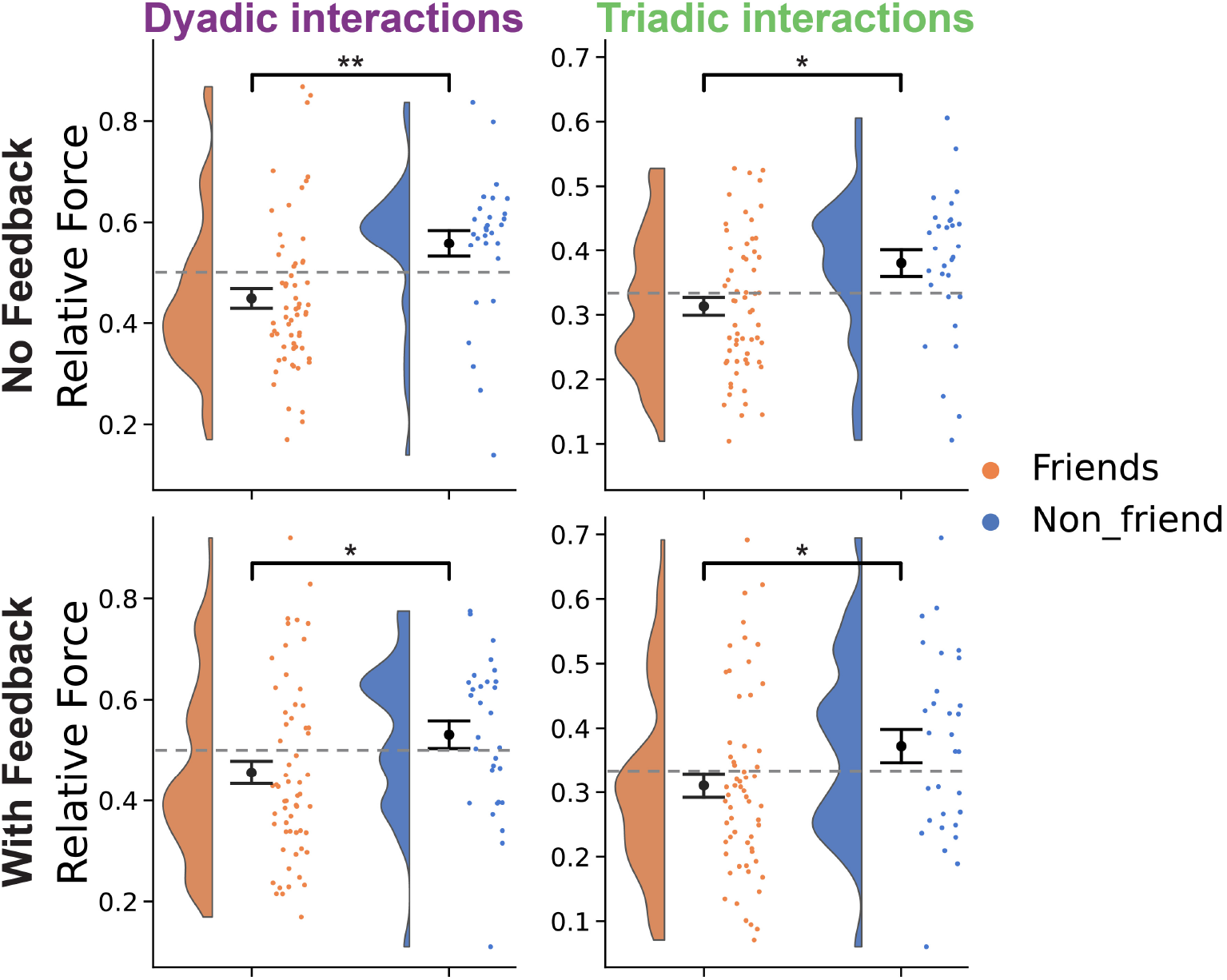
Non-friends produced greater physical force as compared to friends. Non-friends contributed greater physical force to reach the targets in both the dyadic and triadic joint tasks, with and without continuous visual feedback. The striped line corresponds to equal force contribution from all participants. P-values reflect two-sided Wald tests of the corresponding fixed effects from a linear mixed-effects model, after FDR correction. ^∗^*p <* 0.05, ^∗∗^*p <* 0.01.

To unpack how this asymmetry unfolded over time, we next estimated the first derivative of the behavioural force time series, commonly known as rate of force development (RFD) (Maffiuletti et al., 2016), for each trial (Supplementary Figure S2). The peak RFD was then averaged over trials (Figure 3), as a proxy of movement vigour (Marinovic et al., 2017). A linear mixed-effects model showed no three-way interaction between friendship status, task group size, and feedback (*p* ≥ 0.44); however, two-way interactions between friendship status and dyadic trials (*β* = 6.433, 95% CI [3.494, 9.373], SE = 1.500, *z* = 4.29, *p* = 1.79 *×* 10^−5^) and between friendship status and triadic trials (*β* = 10.317, 95% CI [6.915, 13.718], SE = 1.736, *z* = 5.944, *p* = 2.78*×*10^−9^) were observed. Pairwise contrasts showed that non-friends had higher peak RFD in interactive dyadic and triadic trials (Dyadic wFb: *β* = 12.685, 95% CI [3.563, 21.806], SE = 4.654, *z* = 2.73, *p* = 0.010; Triadic wFb: *β* = 14.897, 95% CI [5.616, 24.178], SE = 4.735, *z* = 3.15, *p* = 0.005; Dyadic nFb: *β* = 14.074, 95% CI [4.952, 23.195], SE = 4.654, *z* = 3.02, *p* = 0.005; Triadic nFb: *β* = 17.957, 95% CI [8.676, 27.238], SE = 4.735, *z* = 3.79, *p* = 0.001; FDR corrected across all conditions). No effect of friendship was observed in solo trials (all *p* ≥ 0.13; FDR corrected). This indicates that non-friends produced force more vigorously than friends, specifically during the group coordination tasks.

**Fig. 3.**
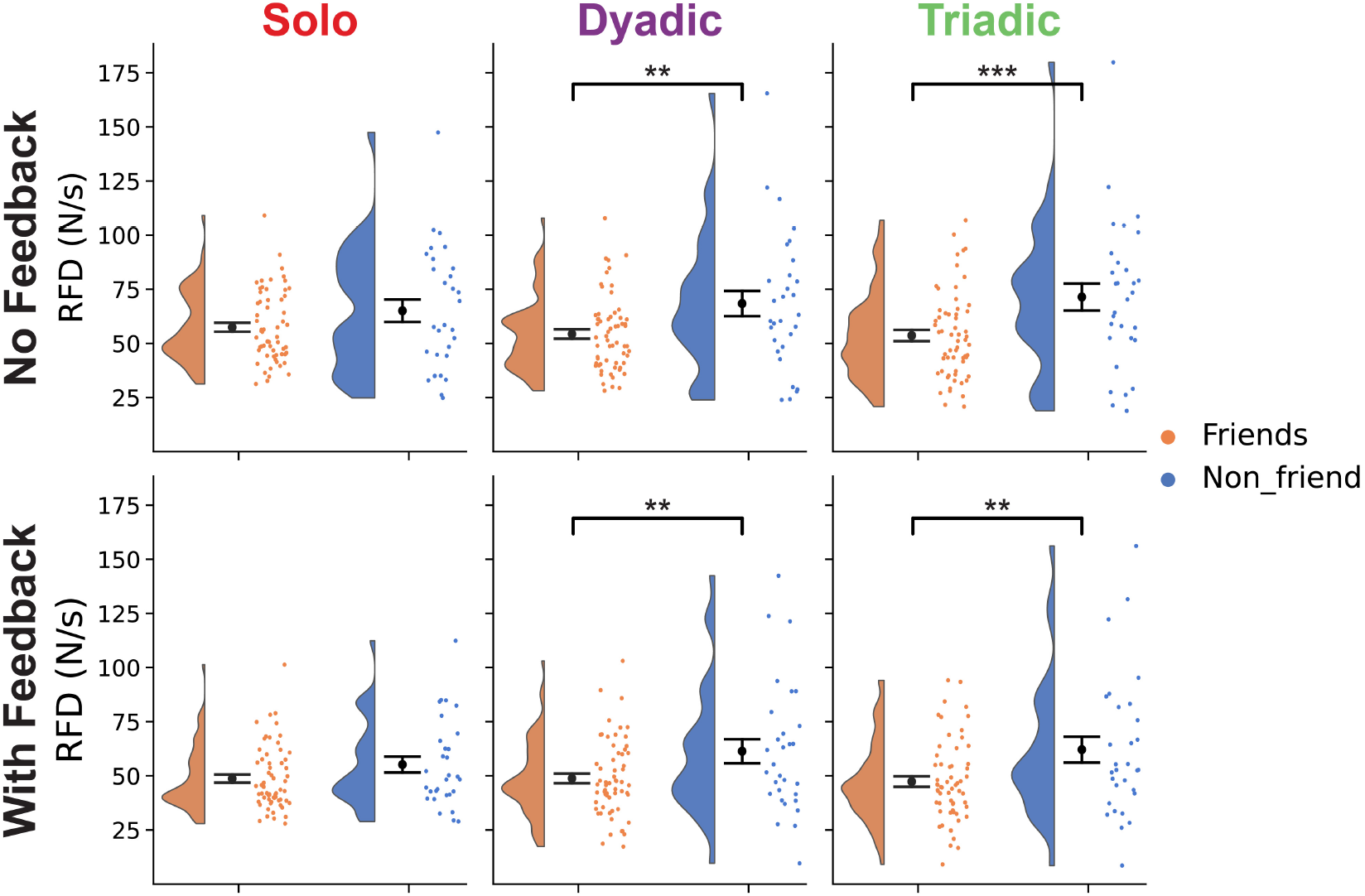
Non-friends produced force more vigorously during interactive joint tasks compared to friends. Non-friends produced force with more vigour, as evident by higher peak RFD compared to friends in interactive dyadic and triadic trials. P-values reflect two-sided Wald tests of the corresponding fixed effects from a linear mixed-effects model, after FDR correction. ^∗∗^*p <* 0.01, ^∗∗∗^*p <* 0.001. RFD: rate of force development.

### 2.3 Force asymmetry was primarily observed in smaller study line classes

We hypothesized that the force asymmetry between friends and non-friends may be explained by the social asymmetries, manifested both as a group-level friendship asymmetry and individual-level outdegree asymmetry, which could be associated with different levels of sensitivity to the social context (Baek et al., 2025). Additionally, social sensitivity could also be influenced by prior familiarity and the size of the class networks, with smaller classes having a higher probability of the students knowing each other. Thus, we investigated the effect of study line class size and number of friends on the force contributions.

Strikingly, a clear association between the size of the study line classes and amount of relative force was observed, with greater force asymmetry observed in smaller classes. A linear mixed-effects model revealed no four-way interaction between class size, friendship status, task group size and feedback (*p* = 0.159); however, two significant three-way interactions were found between class size, friendship status and task group size (*β* = 0.001, 95% CI [0.001, 0.001], SE = 1.96 *×* 10^−4^, *z* = 4.74, *p* = 2.18 *×* 10^−6^) and between class size, friendship status and feedback (*β* = −0.001, 95% CI [-0.001, -0.001], SE = 1.57 *×* 10^−4^, *z* = −6.58, *p* = 4.71 *×* 10^−11^). Simple pairwise analyses showed that in trials with continuous visual feedback, a positive association between the relative force and class size was found for friends, and a negative association for non-friends (all *p* ≤ 0.025; FDR corrected), with trends in the same direction in trials without continuous visual feedback (Figure 4). This shows that the smaller the class size, the more relative force the non-friends contributed in relation to the friends.

**Fig. 4.**
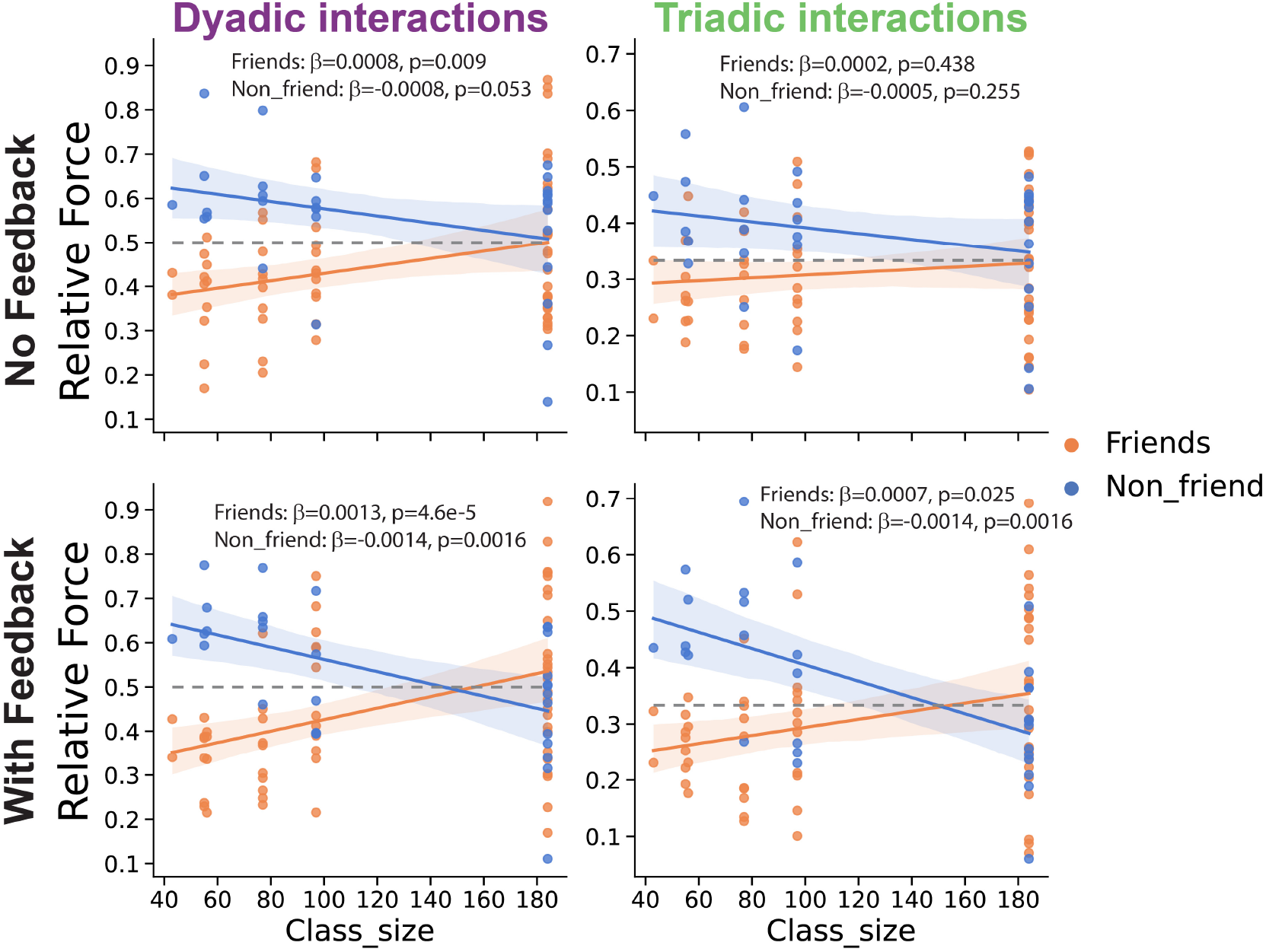
Greater force asymmetry was observed in smaller study line classes. The effect of greater physical force produced by non-friends, and corresponding lower force by friends, was primarily observed for students recruited from smaller study line classes. The striped line corresponds to equal force contribution from all participants. Reported *β* values correspond to estimated slopes from a linear mixed-effects model and p-values reflect two-sided Wald tests of the corresponding fixed effects, after FDR correction.

Similar negative associations in non-friends were also observed between RFD and class size, with significant fourway interactions between class size, friendship status, group size and feedback (Dyadic: *β* = −0.077, 95% CI [-0.152, -0.003], SE = 0.038, *z* = −2.03, *p* = 0.043; Triadic: *β* = −0.141, 95% CI [-0.228, -0.055], SE = 0.044, *z* = −3.20, *p* = 0.001). The simple pairwise contrasts showed no effect among friends, whereas a negative trend was observed for non-friends in interactive trials, although most of these effects did not survive FDR correction (Supplementary Figure S3; uncorrected *p* ≤ 0.041; FDR corrected *p* ≤ 0.098 for non-friends in interactive trials).

We also tested for a direct linear association between the relative force and the number of friends and close friends both inside the classroom and in general, but did not find any reliable linear association after FDR correction (Supplementary Figure S4; all *p* ≥ 0.175; FDR corrected).

### 2.4 Non-friends exhibit increased alpha and beta event-related desynchronization

To investigate the neural mechanisms underlying social coordination, we analysed alpha and beta event-related desynchronization, EEG markers of sensorimotor prediction, preparation, execution, and attention. The Force Game was designed with 30 repeated trials for each condition, which enabled us to increase the signal-to-noise ratio by averaging over trials to perform event-related power analysis across classical EEG frequency bands (Supplementary Figure S5).

Across all conditions, we observed the expected sensori-motor alpha and beta event-related desynchronization (ERD) over left-hemispheric central electrodes contralateral to the moving hand, with desynchronization occurring shortly after the onset of the trial, indicated by the visual appearance of the target force (Figure 5A). This pattern is consistent with classical findings linking alpha and beta desynchronization to motor preparation and execution during voluntary movement (Salmelin and Hari, 1994; Pfurtscheller and Lopes da Silva, 1999; Fujioka et al., 2012; Ross et al., 2017; Zamm et al., 2021).

**Fig. 5.**
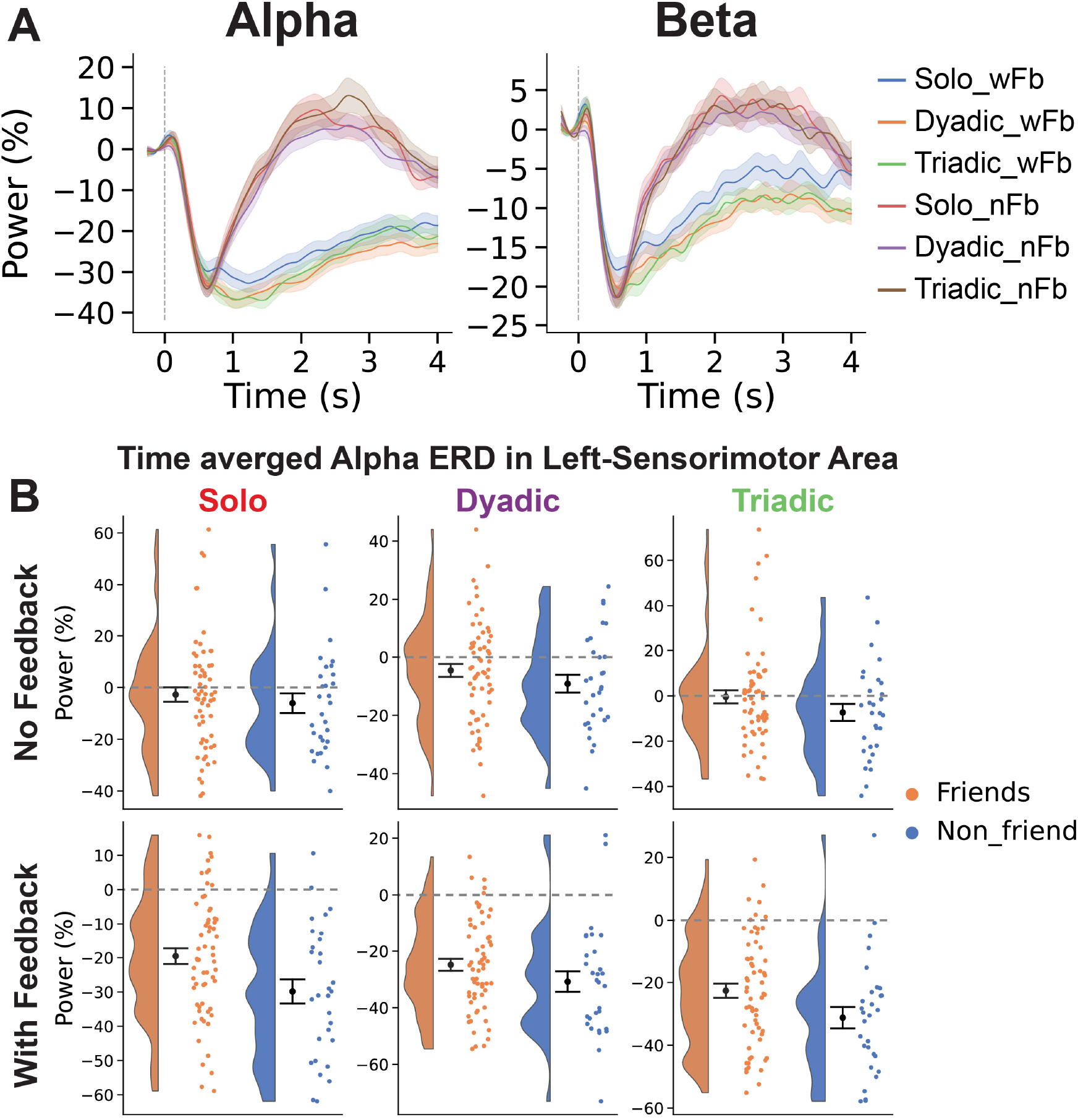
Non-friends exhibit increased alpha/beta event-related desynchronization. A) Example plots of event-related power from electrodes above left sensorimotor area in alpha and beta frequency bands, color-coded based on the different interaction and feedback conditions. B) Time averaged event-related alpha power (0 to 4 seconds) from the left sensorimotor area, plotted separately for friends and non-friends. wFb: with feedback; nFb: no feedback, ERD: event-related desynchronization.

We tested whether these neural responses differed as a function of friendship, i.e., whether non-friends exhibited larger desynchronization during interactive trials. When contrasting friends and non-friends, non-friends exhibited greater ERDs across all frequency bands, illustrated for alpha in Figure 5B. To account for multiple comparisons across time, space (electrodes), and frequency, we conducted non-parametric cluster-based permutation tests (Maris and Oostenveld, 2007; Pernet et al., 2015; Sassenhagen and Draschkow, 2019). In trials with continuous visual feedback, non-friends had significantly greater ERDs across solo (*p* = 0.006), dyadic (*p* = 0.033) and triadic (*p* = 0.01) conditions (Supplementary Figure S6D-F). The clusters consistently spanned approximately 1.1s to 1.7s post-stimulus onset and extended across delta, theta, alpha and beta frequency ranges, and across frontal, lateral-central and posterior regions. No significant differences in ERDs were observed between friends and non-friends in trials without continuous visual feedback (Supplementary Figure S6A-C). There were also no correlations observed between averaged alpha and beta ERD and class network sizes (Supplementary Figure S7 and S8).

Alpha and beta oscillatory activity have been commonly linked to top-down modulation of sensorimotor processing and are thought to play an important role in regulating attention, as they are often modulated by task demand and cognitive effort (Klimesch, 1999; Pfurtscheller and Lopes da Silva, 1999; Mathewson et al., 2012; Keehn et al., 2017; Stecher et al., 2025). Consistent with this interpretation, we observed sustained alpha and beta ERD in trials with continuous visual feedback, whereas power rebounded toward the baseline approximately 2s after trial onset in trials without continuous feedback, where participants made an initial prediction of the required force but in the absence of feedback did not need to adjust this prediction across the trial (Figure 5A). The sustained ERD may thus reflect ongoing attentional demands required to continuously monitor and adapt to visual feedback, consistent with previous findings reporting greater alpha ERD when dyads have visual contact and hence feedback of each other’s movements (Tognoli et al., 2007; Zimmermann et al., 2022). The increased sustained ERD during continuous visual feedback trials was highly significant across the whole brain in solo, dyadic and triadic conditions compared with trials without continuous feedback (Supplementary Figure S9; *p* ≤ 0.001), with clusters consistently spanning approximately 0.7s to 3s in theta, alpha and beta frequency ranges.

Finally, we examined whether ERDs differed between interactive and non-interactive conditions. Alpha and beta ERD were stronger during interactive trials with visual feedback compared to solo trials (Supplementary Figure S10; Dyadic wFb: *p* = 0.002, Triadic wFb: *p* = 0.017; Dyadic nFb: *p* = 0.016, Triadic nFb: *p* = 0.51), consistent with previous work reporting greater frontal and sensorimotor alpha suppression during social interaction (Tognoli et al., 2007; Perry et al., 2011; Naeem et al., 2012; Konvalinka et al., 2014; Zimmermann et al., 2022). One potential explanation is that interaction increases attentional demands by requiring participants to predict and monitor others’ actions in addition to their own (Demos and Palmer, 2023) in order to adapt during coordination.

Taken together, these results indicate that non-friends allocate more attention to the task, particularly when continuous feedback requires ongoing monitoring and adaptation.

### 2.5 Non-friends exhibit increased inter-subject stimulus-locked neural alignment

As ERDs have been associated with multiple neural processes beyond attention, we additionally used inter-subject neural alignment as a proxy for measuring attentional engagement (Dmochowski et al., 2012, 2014; Cohen and Parra, 2016; Madsen and Parra, 2022), to provide converging evidence regarding whether non-friends were more engaged during the task. Correlated components analysis (CorrCA, Parra et al. (2019)) was used to extract neural components that are maximally similar across multiple individuals and enables computation of inter-subject correlation (ISC) during stimulus-locked processing, which has been proposed as a marker of attentional engagement (Dmochowski et al., 2012, 2014; Poulsen et al., 2017; Madsen and Parra, 2022).

Applying CorrCA to event-related alpha and beta power revealed three components with ISC values greater than chance level in both frequency bands (Figure 6A for alpha and S11A for beta). For the first leading correlated component (C1), i.e., the component with the greatest ISC across participants, non-friends exhibited greater ISC compared to friends in trials with continuous feedback across both alpha (Figure 6B; Solo: Δ*M* = 0.110, *d* = 0.622, *p* = 0.014, Dyadic: Δ*M* = 0.082, *d* = 0.572, *p* = 0.015, Triadic: Δ*M* = 0.121, *d* = 0.924, *p* = 0.001; FDR corrected) and beta frequency bands (Supplementary Figure S11B; Solo: Δ*M* = 0.147, *d* = 1.107, *p* = 0.001, Dyadic: Δ*M* = 0.106, *d* = 0.867, *p* = 0.001, Triadic: Δ*M* = 0.093, *d* = 0.687, *p* = 0.002; FDR corrected). The associated topographic distribution for the leading alpha component showed global activation across all channels, with slightly higher weights across lateral visual areas (Figure 6A), whereas the leading beta component had higher weights around left sensorimotor area in addition to occipital areas (Supplementary Figure S11A). In trials without continuous visual feedback, the non-friends only had significantly greater alpha ISC compared to friends in the solo condition (Solo: Δ*M* = 0.096, *d* = 0.58, *p* = 0.015, Dyadic: Δ*M* = 0.043, *d* = 0.311, *p* = 0.167, Triadic: Δ*M* = 0.051, *d* = 0.324, *p* = 0.167; FDR corrected), and no difference was observed in beta ISC (lowest FDR corrected *p* = 0.631). There were also no correlations observed between alpha and beta ISC for the leading component and class network sizes (Supplementary Figures S12 and S13).

**Fig. 6.**
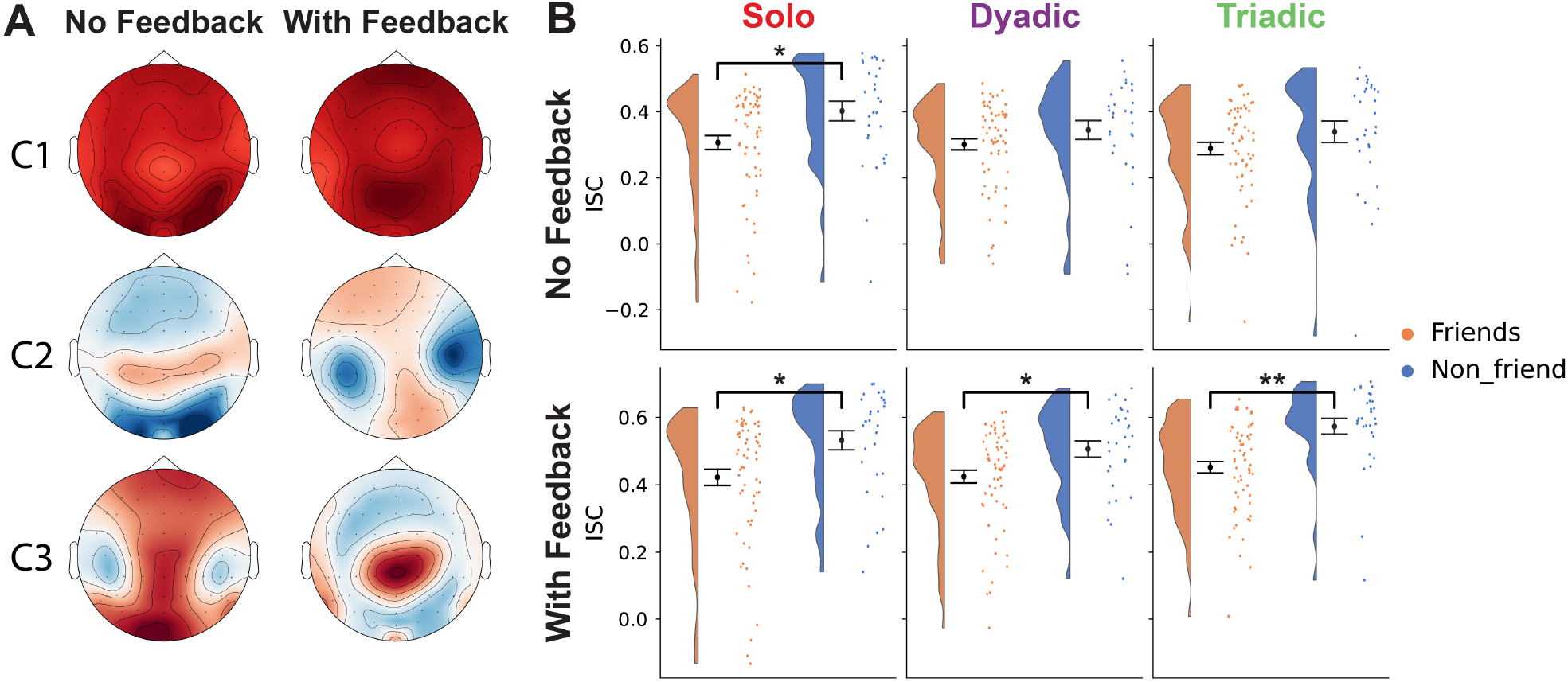
Non-friends exhibit increased inter-subject correlations. A) Three correlated components fitted to alpha event-related power had ISCs above chance levels (*p <* 0.05 estimated with circular shifted surrogate data). B) For the leading component C1, which had the highest ISC, non-friends exhibited greater ISC compared to friends in solo trials without continuous visual feedback and across all conditions with feedback. The topographic maps correspond to projections of the forward model weights, where the polarity is normalized so the Cz electrode weight is positive. ^∗^*p <* 0.05, ^∗∗^*p <* 0.01. P-values were derived using non-parametric permutation tests with FDR multiple test correction. ISC: inter-subject correlation.

To exclude the possibility that higher ISC in non-friends was trivially driven by larger ERD magnitudes, we repeated the CorrCA analysis on individual-level standardized event-related power. The friendship related ISC differences persisted under this normalization, primarily during trials with continuous visual feedback (Supplementary Figure S14), indicating that increased ISC reflects greater temporal alignment rather than differences in signal amplitude (Zimmermann et al., 2024).

Finally, we also estimated inter-subject group cluster phase synchrony (Richardson et al., 2012), a metric complementary to ISC that quantifies neural alignment in the phase domain. Unlike ISC, which is based on correlations and is therefore sensitive to both amplitude and temporal fluctuations, group cluster phase synchrony isolates the consistency of oscillatory phase across individuals. This analysis also revealed greater alpha and beta group phase synchrony in non-friends compared to friends (Supplementary Figures S15 and S16, *p* ≤ 0.0001 for all conditions), indicating that non-friends had greater phase-locked neural dynamics. The increased neural ISC and group phase synchrony in non-friends compared to friends indicates the non-friends had more stimulus-locked neural responses. Based on previous experimental and theoretical research (Hasson et al., 2004; Dmochowski et al., 2012, 2014; Cohen et al., 2018; Madsen and Parra, 2022; Rai et al., 2025), this suggests greater attentional engagement in non-friends performing the Force Game.

## 3 Discussion

Human interaction depends on the ability to coordinate actions while simultaneously inferring others’ behaviour, intentions, and levels of commitment (Sebanz et al., 2006; Michael et al., 2016; Vesper et al., 2017; Sebanz and Knoblich, 2021). Yet little is known about how friendship between interacting individuals influences these moment-to-moment coordination processes. Here, we show that friendship asymmetry within real-world social networks modulates the behavioural and neural dynamics underlying group motor coordination. Across interacting triads consisting of two close friends and one non-friend, we found that non-friends consistently produced greater and more vigorous physical forces than friends. In addition, non-friends exhibited greater alpha and beta ERD, higher neural ISC, and greater group phase synchrony, neural markers associated with action-related and attentional processes.

One potential interpretation of the increased physical effort observed in non-friends is that they occupy a more uncertain social position within the group. Unlike the friends, whose mutual relationship affords greater familiarity and predictability, non-friends must infer both their functional role in the task and their relational standing relative to the others. In socially asymmetric group settings, this uncertainty may motivate individuals to invest more effort as a means of signalling commitment, competence, or value to the group (Leary and Kowalski, 1990; Baumeister and Leary, 1995; Holt-Lunstad et al., 2007; Maner et al., 2007; Pardede and Kovač, 2025). From this perspective, increased force production may serve as a compensatory strategy aimed at fostering affiliation or strengthening one’s position within the interaction (Baumeister and Leary, 1995; Krishna and Götz, 2024). This interpretation is consistent with the increased motor vigour exhibited by non-friends. Movement vigour has previously been linked to reward, motivational states, and commitment (Desmurget and Turner, 2010; Thura and Cisek, 2017; Marinovic et al., 2017; Shadmehr et al., 2019; Marbaker et al., 2025), and the elevated vigour in the absence of external incentives is consistent with greater motivation to engage in the joint task.

Converging neural evidence also supported the interpretation that non-friends exhibited heightened task-engagement. Non-friends showed stronger alpha and beta ERD - proposed neural markers of attention, motor preparation, execution, and prediction (Klimesch, 1999; Pfurtscheller and Lopes da Silva, 1999; Buschman and Miller, 2007; Klimesch et al., 2007; Klimesch, 2012; Kilavik et al., 2013; Stecher et al., 2025) - particularly during conditions requiring continuous monitoring and adaptation to visual feedback of the group force. ERD was also greater during interactive conditions compared to solo conditions, consistent with previous findings showing that social interaction recruits more motor and attentional resources (Perry et al., 2011; Konvalinka et al., 2014; Zimmermann et al., 2022). These effects were accompanied by higher stimulus-locked ISC and greater group cluster phase synchrony (Richardson et al., 2012), suggesting more task-aligned neural processing in non-friends (Dmochowski et al., 2012, 2014; Cohen and Parra, 2016; Parra et al., 2019; Lambrechts et al., 2025). Together, these neural signatures are consistent with heightened attentional engagement with the task and shared feedback signal among non-friends relative to friends. However, these measures cannot distinguish whether the increased task engagement reflects heightened engagement in non-friends, reduced task engagement in friends, or both.

Interestingly, the behavioural and neural findings were differentially related to the broader social environment in which participants were embedded. Specifically, force asymmetry between friends and non-friends was strongest among participants from smaller study line class social networks, whereas no comparable associations were observed for the neural measures. This dissociation suggests that the behavioural and neural effects may reflect partially distinct consequences of social context. One possibility is that broader social-network structure modulates how friendship asymmetries influence behavioural contributions during coordination. In smaller and more tightly connected social environments, where individuals are more visible to one another and interactions more familiar and predictable, friendship asymmetries may exert a stronger influence on how individuals allocate physical effort during coordination, potentially modulated by differences in sensitivity to evaluation, belonging, and inter-personal expectations (Zajonc, 1965; Tetlock and Manstead, 1985; Baumeister and Leary, 1995; Karau and Wilhau, 2020). In contrast, the neural effects may be less sensitive to the broader social network and instead primarily reflect attentional processes associated with the group-level relational asymmetry, with the non-friend occupying a minority position within triads. These possibilities remain speculative, however, as these social and motivational factors were not directly measured.

Additionally, the neural effects were confined to trials with continuous visual feedback, despite the behavioural effects of friendship being present across feedback conditions. This dissociation further suggests that relational context primarily modulates neural processing during phases of interaction that require fine-grained, ongoing sensorimotor adaptation. The temporal profile of the ERD clusters also corresponds to the period following the initial force ramp-up, during which participants actively fine-tuned their force contributions in response to the evolving joint force. In contrast, when continuous feedback is unavailable, coordination relies entirely on internally generated predictions, with no opportunity for online correction, potentially reducing the extent to which social and relational factors can modulate the ongoing coordination. Thus, these findings indicate that continuous adaptation rather than internal prediction determine when relational effects are expressed in neural dynamics. Altogether, these findings suggest that the impact of friendship is not uniform, but depends on both online coordination mechanisms and the broader social context the participants are embedded within.

Several interpretations for the role of friendship remain possible. Increased force contribution may reflect a compensatory response to social asymmetry, whereby non-friends feel a greater need to prove themselves. Alternatively, force asymmetry may be attributed to the fact that friends allocate greater attentional resources toward one another, rather than task-related information, hence producing force more hesitantly and with less vigour in interactive trials. In order to disentangle the underlying mechanisms, future studies should incorporate subjective measures of the need to belong and social motivation.

The observed effects of friendship are unlikely to be explained by low-level baseline motor or electrophysiological factors. No individual differences in baseline maximum physical strength, maximum motor vigour (Supplementary Figure S17), response times (Supplementary Figure S18) and behavioural performance in solo tasks (Supplementary Figure S19) were observed between friends and non-friends. Additionally, performance in dyadic friend – friend interactions was also on par with friend – non-friend interactions (Supplementary Figure S20). Moreover, the greater alpha and beta ERD in non-friends was not attributable to the greater force produced, as ERD magnitude was not associated with the amount of produced force (Supplementary Figures S21 and S22). These findings suggest that the observed behavioural and neural differences reflect higher-order cognitive processes related to social context, rather than differences in physical capacity or movement execution.

While the present findings provide insight into how friendship influences coordination, several limitations remain. Although the paradigm offers a controlled means of quantifying joint motor coordination, it captures only one facet of social group behaviour, and the influence of social asymmetry may differ in tasks involving other social, cognitive, or motivational demands (Zajonc, 1965; Karau and Williams, 1993), such as face-to-face verbal interactions (Sievers et al., 2024). The specificity of our task could also explain why our finding of greater neural similarity among non-friends appears to contrast with prior social neuroscience results showing that friends tend to exhibit more similar neural activity at rest (Hyon et al., 2020b) or during passive video watching (Parkinson et al., 2018; Hyon et al., 2020a; Shen et al., 2025). In the present study, neural similarity reflects stimulus-locked, task-evoked processing during active coordination rather than intrinsic similarity in resting-state activity or passive perception. Under these conditions, stronger ISC in non-friends likely reflects greater alignment to the shared task dynamics, rather than stable overlap in neural representations.

In addition, the analysis of class-level social networks was constrained by incomplete network coverage, limiting estimation of higher-order network metrics. On average 66.4% of the students reported their connections, precluding reliable estimation of network parameters such as eigenvector centrality, density, clustering, or assortativity (Kossinets, 2006; Smith and Moody, 2013; Marks et al., 2013). Consequently, we focused on outdegree which can be more robustly estimated under partial sampling. Non-friends also reported fewer friends overall compared to the participants recruited as friends, meaning that friendship status within the triad was not entirely independent of broader individual social connectedness. Although the number of friends did not reliably predict relative force after correction for multiple comparisons, future studies should more directly dissociate an individual’s friendship position within the interaction group from their broader network position.

In summary, our findings demonstrate that differences in social relationship - friend versus non-friend - systematically modulate how individuals allocate physical effort and engage with a shared task during group coordination. Non-friends contributed more force and acted with greater vigour, accompanied by neural dynamics consistent with heightened task engagement. Friends, by contrast, showed behavioural and neural mechanisms consistent with reduced vigilance and lowered task-related engagement in relation to non-friends. Together, these findings show how social relationships within real-world social groups are dynamically embodied in the behavioural and neural processes that unfold during real-time collective action.

## 4 Methods

### 4.1 Social network mapping

378 students across seven bachelor program study lines (range: 43-184 students, *M* = 83.14, *SD* = 44.26) at the Technical University of Denmark (DTU), gave informed consent to have their social networks in their study line programs mapped. Specifically, 28 to 113 students per study line filled our questionnaire (*M* = 54.0, *SD* = 27.09), corresponding to a response rate ranging from 50.6% to 95.7% (*M* = 66.4%, *SD* = 13.05%) of the size of the study lines.

In practice, we gave a brief 5 min presentation physically in the classrooms about the project and experiments, and asked students to fill out questionnaires in which they checked off the fellow students they considered a friend, close friend, or leave it blank to indicate the fellow student was not a friend (Figure 1A). Explicit definitions of friends and close friends were provided (see Supplementary Methods for details). Participants recruited for the experiment were also asked to fill an additional questionnaire about their friends and close friends outside their study programs, divided into workplace, hobbies, social media, studies and “other” categories to promote recall, upon arriving in the lab.

### 4.2 Experimental participants

From the class social networks, 93 healthy right-handed participants (35 females) between 18 and 35 years (*M* = 21.4, *SD* = 2.29) were recruited in 31 triads (19 mixed-sex, 12 same-sex, including 3 all-female, 9 all-male triads) composed of two reciprocal close friends and one non-friend. The study was conducted according to the Declaration of Helsinki and were approved by the Scientific Ethics Committee for the Capital Region of Denmark (H-23013244). Participants in the experimental task were financially compensated for their participation.

### 4.3 The Force Game paradigm

The three participants sat facing away from each other in a triangular formation (Figure 1B). Each participant had a screen and was equipped with a custom-made 3D printed pressure measuring (force) device, which we calibrated using a set of weights to estimate the forces in Newtons (Supplementary Figure S23). The participants were instructed to hold the force devices with their right hand and press with their right thumbs to reach a target force, either individually or jointly. Practice trials were also performed prior to the experiment, to familiarize the participants with the devices. The experiment consisted of 300 trials, across 10 conditions with varying group sizes (solo, dyadic and triadic) and feedback. We manipulated whether the interaction recruited adaptive or predictive mechanisms, by either giving them continuous visual feedback, or no feedback across the 4-second trial, followed by discrete feedback of the reached force at the end of the trial (see Supplementary Methods for details). While performing the Force Game, three-person EEG was simultaneously recorded. The experiment was designed using Psychopy (Peirce et al., 2019) and took around 45 min, and the whole procedure including EEG setup took around 1 hour and 30 min.

### 4.4 Data acquisition

To synchronize the force devices, we stored the force measurements corresponding to when the screens updated the visual display. This resulted in the behavioural force data being acquired at 25 Hz, due to computational limitations associated with dynamically updating across three screens simultaneously from one PC (see Supplementary Methods for details). Nonetheless, visual inspection confirmed that 25 Hz was enough for the stimuli to visually appear continuous, and 25 Hz was enough to capture smooth force trajectories.

EEG was recorded using three synchronized, daisy-chained 64-channel Biosemi (Amsterdam, The Netherlands) ActiveTwo EEG set-ups in a 10–20 system, at a sampling frequency of 2048 Hz. EEG caps with 64 channels were positioned such that Cz was centered on the head, at the mid-point between the nasion and inion, and the left and right ear. Conductive gel (SignaGel Electrode Gel, Parker Laboratories, Farfield, NJ) was used to improve signal reception, and electrodes with offsets above *±* 20 mV were adjusted in order to keep them as close to 0 as possible. Any visually bad electrodes at recording were noted down for exclusion during preprocessing.

One participant was excluded from analysis due to non-compliance with the task, e.g. not pressing at all in many trials, resulting in failing to reach the target force in half of solo conditions with feedback, where the average across participants was 98.9%. Specifically, the solo, dyadic and triadic conditions involving the non-compliant participant were discarded, but the solo and dyadic conditions with the remaining two participants in the triad were kept. Thus, the final analysis was performed on the remaining 62 friends and 30 non-friends.

### 4.5 Behavioural analysis

A successful trial was defined as a trial where the force trajectory (solo or group) reached within *±*2N of the target force, and stayed at the target for at least 200 ms. The accuracy (success rate), latency to success, and stability (duration the force stayed around the target) was computed. Relative force was estimated as the force contribution relative to the target force in group tasks at time of success. The instantaneous rate of force development was computed as the first derivative of the force time series. The 10 experimental conditions were collapsed into 6 conditions (group size x feedback). The same 6 collapsed conditions were applied for EEG analysis (see Supplementary Methods for details).

### 4.6 EEG preprocessing

The EEG data were processed using MNE-Python 1.6.1 (Gramfort et al., 2013). First, the data were bandpass filtered at 1–40 Hz (Hamming), downsampled to 500 Hz, and segmented into epochs (-0.5 to 5.5 seconds relative to trial onset). Bad epochs and channels with gross artefacts were rejected by visual inspection. The bad channels were interpolated from adjacent channels using spherical spline interpolation (Perrin et al., 1989) followed by re-referencing to the common average. Ocular, heart and muscle related artefacts were removed using independent component analysis with the number of components set to 32. The automatic detection of muscle-related components in MNE-Python was employed to guide the manual determination of muscle artefacts. Autoreject 0.4.3 (Jas et al., 2017) was employed to catch any remaining artefacts by interpolating specific bad channels within each epoch, followed by a guided final manual visual inspection with highlighted bad epochs determined by the algorithm. Using Autoreject this late in the preprocessing pipeline made it prone to false-positives if the signal was already clean, hence a manual check was performed and no epochs were dropped automatically. On average 3.45 (*SD* = 5.12) number of epochs were dropped, 1.1 (*SD* = 2.4) number of channels were interpolated and 15.66 (*SD* = 4.27) independent components were corrected. The EEG features were estimated in the five canonical frequency bands: delta (1–4 Hz), theta (4–8 Hz), alpha (8–13 Hz), and beta (13–30 Hz).

### 4.7 Event-related power

The time-frequency representation (TFR) of the epoched EEG data was computed using Morlet wavelets. A variable window length that decreased with increased frequency was used to obtain greater temporal resolution at higher frequencies. The power was computed for each frequency band and averaged across epochs for each of the 6 collapsed conditions. Percentage based baseline correction was applied on the trial averaged TFR using the pre-stimulus interval (-250 – 0 ms), i.e. on a single-subject level for each condition to avoid positive bias from performing percentage based baseline correction on a single-trial level (Hu et al., 2014).

### 4.8 Inter-subject correlation

ISC was estimated using correlated component analysis (Parra et al., 2019). Specifically, for each frequency and condition, we applied correlated component analysis on the TFRs, with participants on the first axis to compute stimulus-related power components that maximally correlate between subjects. To determine similar components across group size conditions (but separately for visual feedback), we concatenated the different group conditions along the first axis, in order to make ISC values comparable across group sizes. To determine an effect of friendship, ISC was separately computed among friends and among non-friends, using all-to-all pairwise Pearson’s correlations of the extracted components across all triads, to estimate aligned stimulus-locked neural responses.

### 4.9 Group cluster phase synchrony

Group cluster phase synchrony was estimated following Richardson et al. (2012). Briefly, epochs were averaged to obtain event-related potentials, followed by filtering for each canonical frequency band. Hilbert transform was applied to estimate the analytic signal, and the phase was extracted. Conceptually, group cluster phase synchrony is similar to the classical connectivity metric phase-locking value (Lachaux et al., 1999), except relative phase is not computed between individual electrodes or subjects, but between each individual to the group averaged phase, for each electrode. To determine an effect of friendship, group cluster phase synchrony was separately computed among friends and among non-friends.

### 4.10 Statistical analysis

Results are shown as mean with standard error bars. The shaded areas in the correlation plots correspond to the 95% confidence intervals. Two-sided non-parametric permutation tests (10000 permutations) with FDR correction were used to test for group mean differences. The FDR correction was applied to each metric separately, but across conditions. QQ-plots were assessed to evaluate for normality, and Spearman’s correlation was used to test for significant correlations. For all statistical tests, a p-value *<* 0.05 was considered significant for rejection of the null hypothesis and the number of asterisks corresponded to each of the following significance levels: ^∗^*p <* 0.05, ^∗∗^*p <* 0.01 and ^∗∗∗^*p <* 0.001. Effect sizes were estimated with Cohen’s *d* (see Supplementary Materials for equation).

Behavioural features estimated at the individual-level, i.e. relative force and RFD, were analysed using linear mixed-effects models to account for the hierarchical structure of the data, with repeated trials nested within individuals and individuals nested within dyads or triads, to take into account the non-independence arising from shared task context during collective action. Specifically, we fitted 3-way mixed effects models to investigate the effect of friendship on relative force and RFD of the form: RelativeForce ∼ FriendStatus *×* GroupSize *×* Feedback + (1 | Triad) + (1 | Participant). Here the fixed effects included the main effects of interest, i.e. friend/non-friend status and experimental conditions as well as their interactions. Random intercepts were included for triads and participants to account for the nested structure (see Supplementary Materials for more details). P-values for fixed effects and pairwise contrasts were derived using two-sided Wald z-tests.

Non-parametric cluster-based permutation tests (Maris and Oostenveld, 2007; Sassenhagen and Draschkow, 2019) were applied to handle the multiple comparison problems when taking time, space (electrodes) and frequency into account for estimating group differences in ERDs and group cluster phase synchrony (see Supplementary Methods for details).

## Data and code availability

The code that support the findings of this study are openly available on https://lab.compute.dtu.dk/glia/force-game-eeg-analysis. The dataset are available via the DTU data repository: https://doi.org/10.11583/DTU.31369621.

## Author contributions statement

Q.L. and I.K, conceived and designed the study and experiments. Q.L., A.D., K.K., and I.K. conducted the experiment and collected the data. Q.L. implemented the code. Q.L. and I.K. designed the analysis and generated the first draft of the manuscript. Q.L., A.D., K.K., and I.K. contributed to interpretation of the results and reviewed the manuscript.

## Acknowledgements

This work is supported by the Villum Young Investigator grant (project no. 37525) and the Carlsberg Semper Ardens: Accelerate grant (CF22-1251).

