## Supplementary Materials for "Friendship modulates physical effort and neural dynamics during group coordination in shared social networks"

### Supplementary Methods

#### Social Network Mapping

Explicit definitions of friends and close friends were provided for the social network mapping questionnaires. A friend was defined as *a person whom you like to spend your free time, enjoy informal social activities, such as going for lunch, dinner, drinks, films, etc., and help each other out in a practical way* (Burt, 1995; Spencer and Pahl, 2006; Parkinson et al., 2018; Sievers et al., 2024). A close friend was defined as *a person whom you not only help each other and enjoy each other's company but also share personal information and provide emotional support* (Spencer and Pahl, 2006). The social network surveys were administered during the students' first (2nd semester) or second (3rd or 4th semester) academic year in their bachelor program, as the courses taken by the students' in the early semesters of their bachelor programs are primarily mandatory courses, hence the students have a high overlap of common courses and are more likely to know each other within each study program.

#### Experimental participants

The sample size was estimated based on pilot data. An a priori power analysis, assuming 80% power and a significance threshold of $\alpha = 0.05$, indicated a target sample size of approximately 30 triads. Recruitment proceeded study line by study line, such that once eligible participants within a study line were exhausted, additional classes were mapped.

Gender composition was not used as a recruitment criterion, as simultaneously matching participants on friendship configuration and gender composition would have substantially reduced the pool of eligible participants and made recruitment infeasible in several study lines. Moreover, the triadic design yielded multiple combinations of friendship role and gender composition (e.g., MMM, MFM, FFM, MMF, MFF, FFF, where the final position denotes the non-friend), thus incorporating gender would have resulted in a large number of subgroup combinations with insufficient sample sizes for reliable statistical comparisons. Consequently, the study was not powered to evaluate interactions between gender composition and friendship asymmetry.

#### The Force Game paradigm

The Force Game was conducted inside an electrically shielded room. The three participants sat facing away from each other in a triangular formation with 50 cm between each pair of chairs. The screens were placed 60 cm in front of the center of each chair, and would display the force (either solo or joint) as an expanding white circle (the bigger the force, the larger the circle) and the target force as a hollow blue circle. When the produced force was within $\pm$ 2N of the target force, the whole circle would turn blue indicating the participants successfully reached the target.

The participants were explicitly instructed to reach the target force and subsequently keep the force at the target for the whole duration of the 4-second trial. The radius of the displayed force and target force was based on the log-transformed forces in Newtons, to be visually sensitive to smaller forces. Preliminary pilot runs estimated 50N as a reasonable max force and in order to enable all participants to be able to reach the target and taking fatigue into account, we set the target forces to vary as 8N, 10N, 12N, 14N or 16N (corresponding to around 15 to 30% of max force production). The averaged max force during the practice trials from all 93 participants was 49.59N, consistent with our pilot results.

In the practice trials the participants were instructed to freely press, press as hard as they could, or reach the target force individually or jointly. The experiment consisted of 300 trials, across 10 conditions with varying group sizes (solo, dyadic and triadic) with and without continuous visual feedback. There were three duo tasks, i.e. P1 and P2, P1 and P3 or P2 and P3 interacting while the last participant's force contribution was excluded from the joint force, a manipulation they were not aware of. In group trials, only the averaged collective joint force was shown, and the contribution from each participant was $1/2$ or $1/3$ the actual individually measured force in the dyadic and triadic trials respectively, to take into account the increase in group size. During the experiment, each condition was performed in blocks of five trials, corresponding to the five target forces (in randomized order), followed by a self-paced break. Prior to each 4-second trial, a fixation cross was presented on the screen, with a jittered 1 to 2 seconds duration, randomized for each trial to prevent EEG synchronization due to the predictability of stimulus onset timing. Prior to each block, a condition cue was presented with the text *Individual Task* or *Group Task* followed by a green emoji with a single person or an orange emoji with three people, and the text *Continuous Feedback* or *No Continuous Feedback*. No distinction in condition cue was given for dyadic or triadic trials, and no explicit information about who was excluded in duo trials was provided. For trials with no continuous feedback, only the hollow blue circle corresponding to the target force was shown during the 4-second trial, followed by a static image of the joint force at the end of the trial, shown for 1 second. All 10 conditions were also administered in blocks (in randomized order), repeated six times (Figure 1C), to avoid the same conditions being consecutively performed (on top of the 5 repeated trials). Longer forced breaks of 1 min were also given once 100 or 200 trials had been completed to ensure the participants relaxed their thumbs.

**Force device specifications**

The custom-made force devices send a constant voltage to the Arduino, which converts the analogue voltage signal to a digital 8 bits signal and transfers it to the stimuli PC. The force devices have a pressure-sensitive button, which changes the resistance when pressed and thus the voltage will change accordingly. To calibrate the force devices, we used a set of weights (295g, 500g, 1000g, 2000g and 5000g) to estimate the forces in Newtons (Supplementary Figure S23) using the equation $F = m \times g$, where $g = 9.8m/s^2$ is the standard acceleration corresponding to gravity on Earth. Third-order polynomials with intercepts forced to 0 was fitted to the measured calibration data to obtain the calibration curves for each force device.

Based on the calibrations with constant weights, we observed measurement errors in the range of 0.1N. Each force device sends a continuous stream of numbers (more than 1000/sec), but to synchronize the measurements, we stored the force measurements corresponding to when the screens updated the visual display, which resulted in a sampling rate of 25 Hz. Despite using 60 Hz monitors, it took computational time to visually create the elements and display on the three screens simultaneously from one PC, hence the stimuli update frame rate and force sampling rate was lower than 60 Hz.

**Behavioural analysis**

The behavioural force measurements were linearly interpolated to common timepoints, in fixed 40 ms intervals (25 Hz) starting from trial onset to end of trial at 4 seconds, in order to analyse across trials. The instantaneous rate of force development (RFD) was approximated by taking the difference in force between each sample, divided by the time difference: $RFD = \Delta F/\Delta t$. Coefficient of variability was computed 2 s after trial onset, to avoid the build-up of force when approaching the target, while focusing on the part where participants had to stay around the target. The 10 experimental conditions were collapsed into 6 conditions (3 x 2, corresponding to group size x feedback), by combining the two dyadic conditions where each participant was actively contributing to the joint force, and removing the dyadic condition that each participant was excluded from. The same 6 collapsed conditions were applied for EEG analysis.

**Inter-subject correlation**

To account for the positive bias induced by higher SNR in the TFRs from the collapsed dyadic condition (60 trials as opposed to the 30 trials), we calculated ISC separately for TFRs computed from each of the two uncollapsed included dyadic conditions for each participant, followed by averaging after ISC was computed, as a higher SNR condition can arbitrarily increase inter-brain synchrony estimates (Zimmermann et al., 2024). To ensure the observed difference in ISC was not confounded by a difference in number of subjects for each group (two friends for each non-friend), we performed 1000 permutations where we performed the ISC estimation on 30 randomly sampled friends (without replacement) to create a null distribution and no difference was seen compared to the mean ISC for the original full sample of friends (lowest $p = 0.951$ across alpha/beta ISCs across all conditions). To rule out that the increased ISC in non-friends was driven by increased ERD in non-friends, we also repeated the correlated component analysis on subject-level standardized TFRs and observed similar results.

**Group cluster phase synchrony**

Group cluster phase synchrony was estimated following Richardson et al. (2012). After extracting the phase signal for each electrode, the group cluster-phase time series was estimated as the averaged phase time series across individuals, from which relative phase time series for each individual could be computed, followed by estimation of the degree of synchrony for each individual with respect to the group. Similar to ISC, we accounted for potential positive bias induced by higher SNR from collapsed dyadic conditions by performing the inter-subject analysis separately for each uncollapsed condition, before averaging the dyadic conditions after group synchrony was computed.

**Statistical analysis**

Effect sizes were estimated with Cohen's $d$ computed as:

$$d = \frac{\bar{M}_1 - \bar{M}_2}{s_{\text{pooled}}} \tag{1}$$

where $\bar{M}_1, \bar{M}_2$ are the group means and the pooled standard deviation defined as:

$$s_{\text{pooled}} = \sqrt{\frac{(n_1 - 1)s_1^2 + (n_2 - 1)s_2^2}{n_1 + n_2 - 2}} \tag{2}$$

Besides binary main effects, we also had models with continuous variables, e.g. class size or outdegree: $\mathrm{RFD} \sim \mathrm{ClassSize} \times \mathrm{FriendStatus} \times \mathrm{GroupSize} \times \mathrm{Feedback} + (1 \mid \mathrm{Triad}) + (1 \mid \mathrm{Participant})$. Random slopes for key predictors were explored but resulted in near-zero variance estimates or convergence issues, indicating limited heterogeneity in effects across triads. Therefore, the final linear mixed-effects models included random intercepts only.

1000 permutations were used for the non-parametric cluster permutation tests contrasting ERDs, due to high computational demand with the high dimensionality over time, space and frequency, while 10000 permutations were used for group cluster phase synchrony that only had space and frequency dimensions. A T-statistic with cluster forming threshold of 2 was employed.

### Supplementary Figures

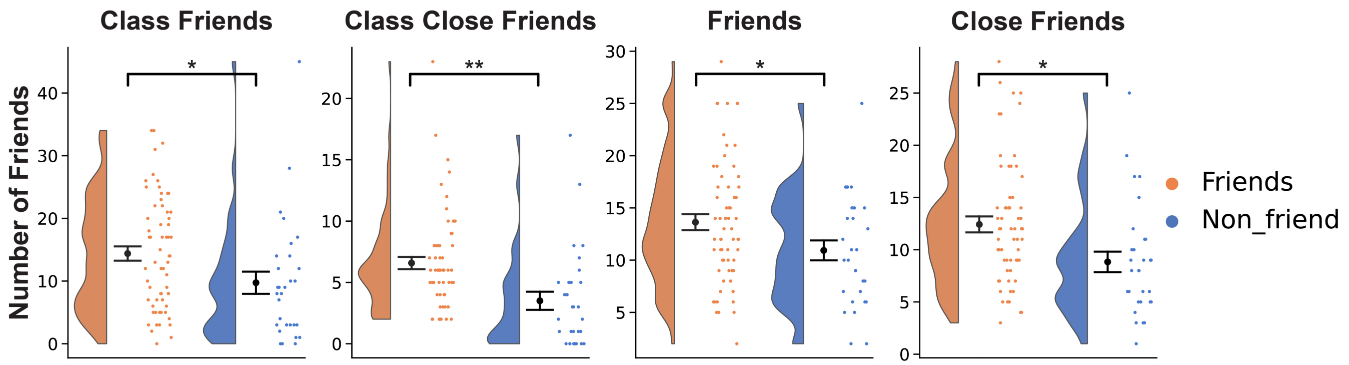


**Fig. S1**. **Non-friends had fewer reported friends and close friends relative to friends.** Participants filled out a questionnaire about whom they were friends and close friends with in their study line classes, and another follow-up questionnaire in the laboratory where they were asked to list friends outside their classrooms to get a complete estimate of their number of friends. Non-friends had fewer friends across all four categories. $^{*}p < 0.05$, $^{**}p < 0.01$. P-values were derived using non-parametric permutation tests with FDR multiple test correction.

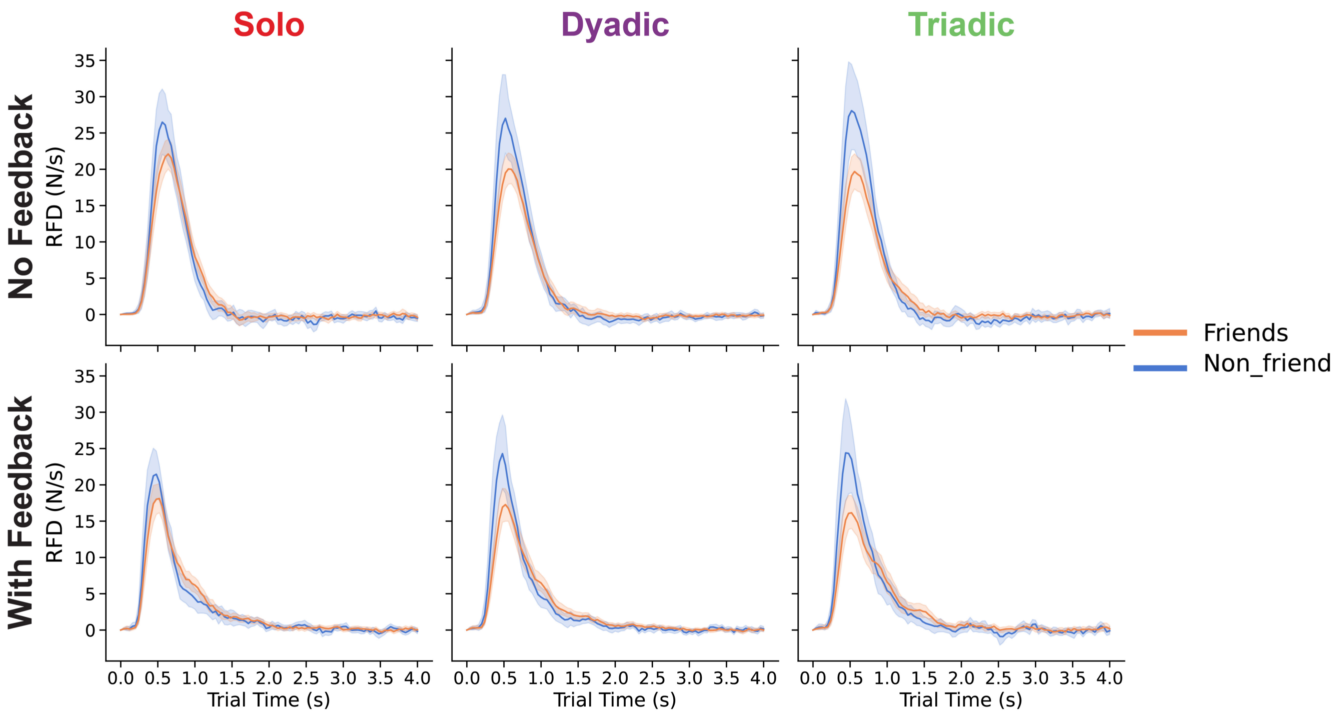


**Fig. S2**. **Rate of force development.** RFD, the first derivative of the force time series, reflects how quickly the force changed at a given time point. A clear peak in RFD was observed when the participants started pressing the force devices, followed by a subsequent decrease as they approached the target. The peak RFD in non-friends was higher than in friends during interactive trials. RFD: rate of force development.

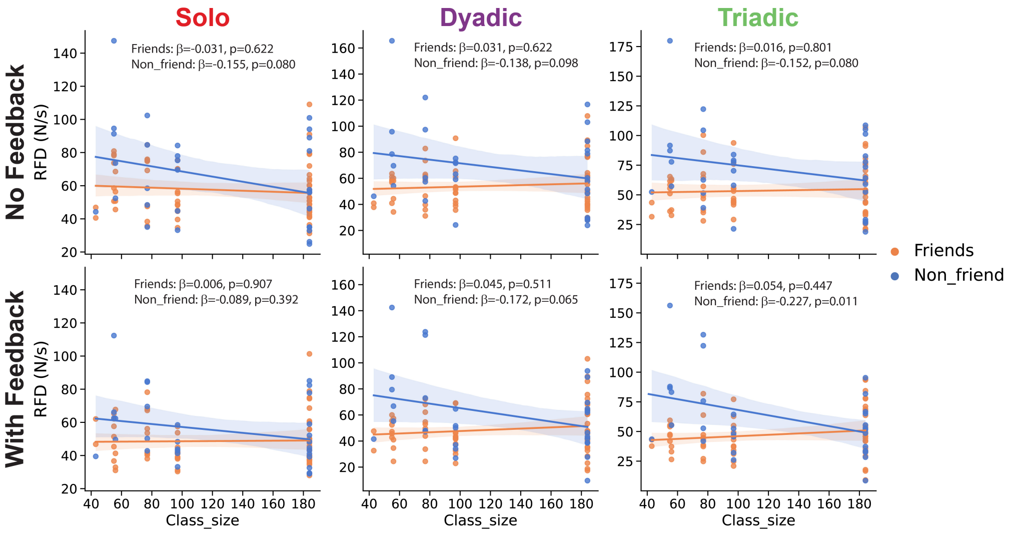


**Fig. S3**. **Correlations between rate of force development and class social network sizes.** A negative trend between RFD and the size of the class social networks were observed for non-friends, however most of the correlations were not significant after FDR correction. Reported $\beta$ values correspond to estimated slopes from a linear mixed-effects model and p-values reflect two-sided Wald tests of the corresponding fixed effects, after FDR correction.

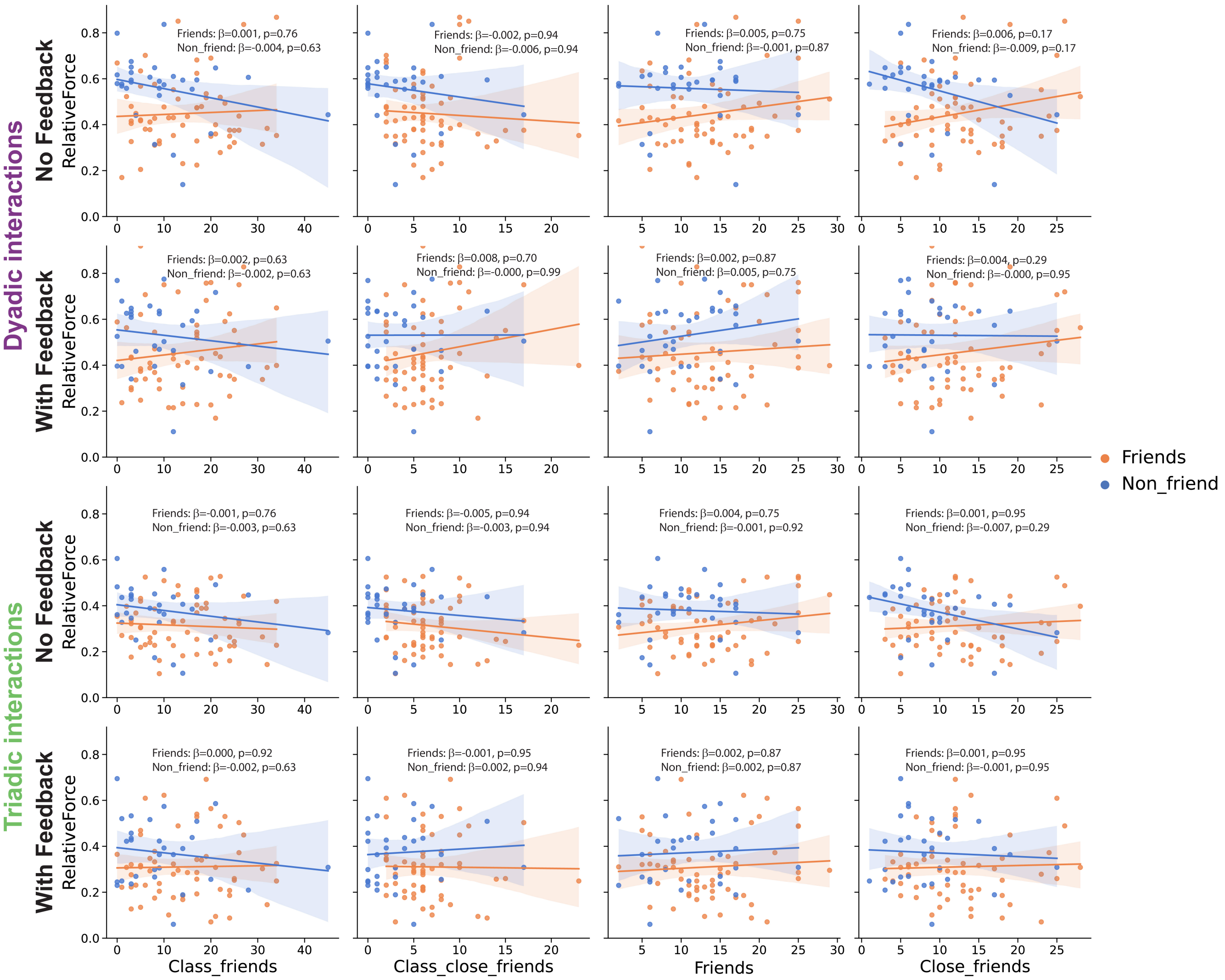


**Fig. S4**. **Correlations between number of friends and relative force.** No correlations were observed between relative force and the number of class friends, close class friends, friends outside classroom and close friends outside classroom. Reported $\beta$ values correspond to estimated slopes from a linear mixed-effects model and p-values reflect two-sided Wald tests of the corresponding fixed effects, after FDR correction.

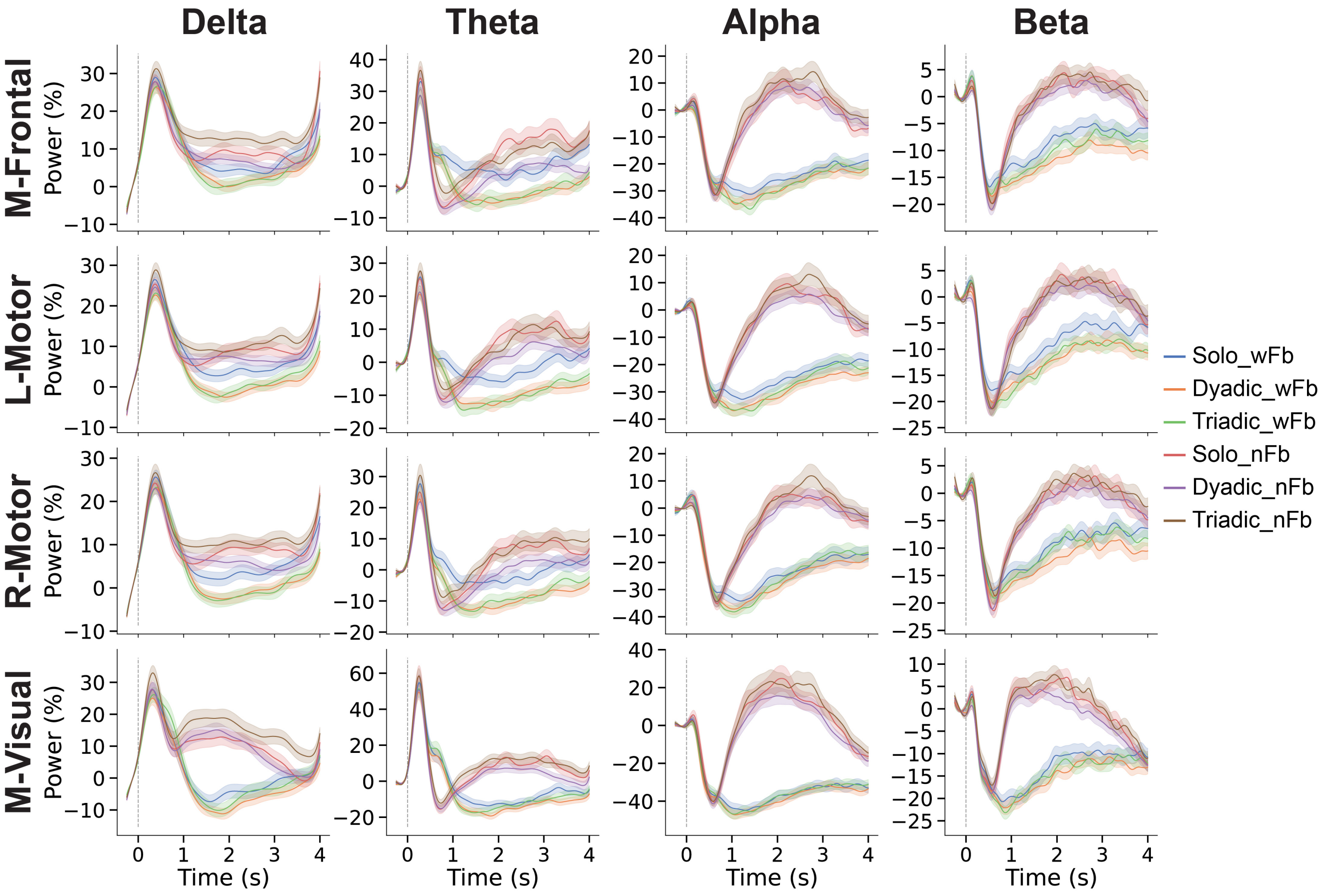


**Fig. S5**. **Event-related power analysis.** Examples of event-related power in delta, theta, alpha and beta frequency bands averaged over electrodes from different brain areas. Mid-Frontal: F1 , Fz, F2, FC1, FCz, FC2, C1, Cz, C2. Left-Motor: FC5, FC3, FC1, C5, C3, C1, CP5, CP3, CP1. Right-Motor: FC6, FC4, FC2, C6, C4, C2, CP6, CP4, CP2. Mid-Visual: PO3, POz, PO4, O1, Oz, O2. wFB: with feedback; nFB: no feedback, ERD: event-related desynchronization.

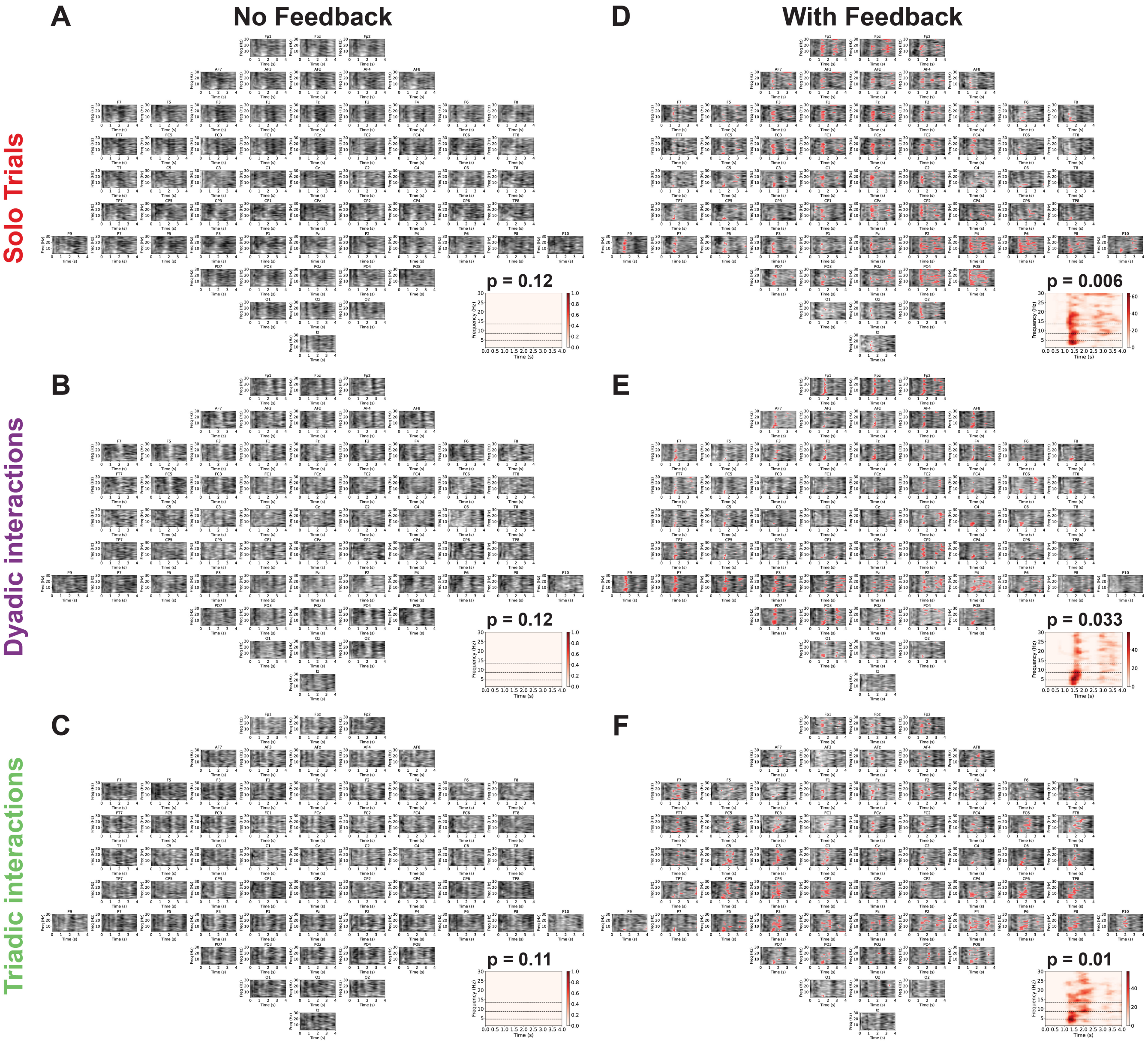


**Fig. S6**. **Non-parametric cluster-based permutation test for effect of friendship on event-related power.** T-statistics are shown in spectrograms with time on the x-axis and freq on the y-axis, with subplots for each electrode, ordered spatially with the top row corresponding to most anterior positions. Darker colors indicate greater t-values, with red colors indicating the data points are part of a significant cluster with more desynchronization in non-friends compared to friends. A-C) No effect of friendship was observed in trials without continuous visual feedback. D-F) In trials with continuous visual feedback, non-friends had significantly lower power (greater desynchronization). Clusters consistently found across solo, dyadic and triadic conditions spanned from approximately 1.1s to 1.7s and across delta, theta, alpha and beta frequency ranges, in frontal, lateral-central and posterior regions. The insets show summary spectrograms collapsed over the spatial dimension (electrodes), counting in how many electrodes, each data point in the spectrograms were part of the significant cluster.

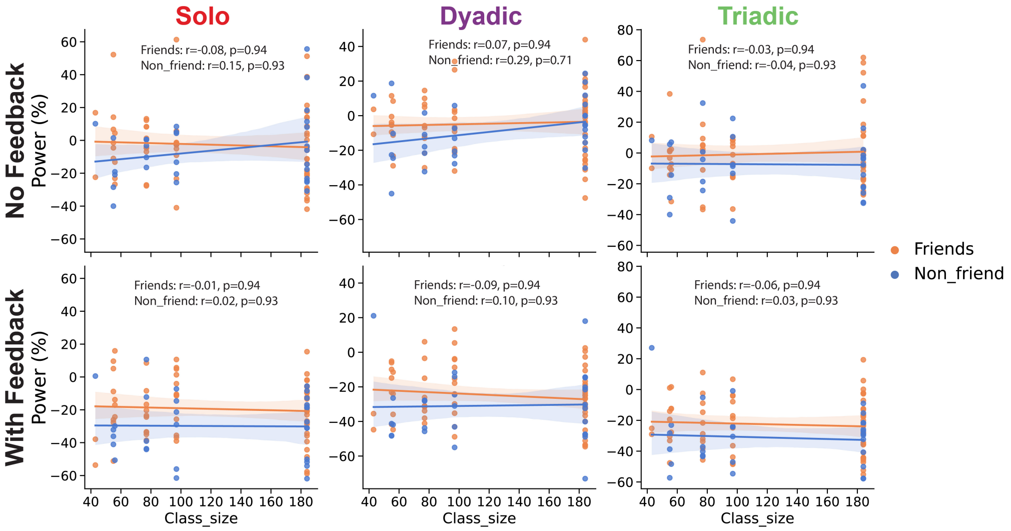


**Fig. S7**. **No correlations between alpha ERD and class social network sizes.** No associations were observed between trial averaged left-sensorimotor alpha ERD and the size of the class social networks. Spearman correlation coefficients were computed and the p-values shown have been FDR corrected.

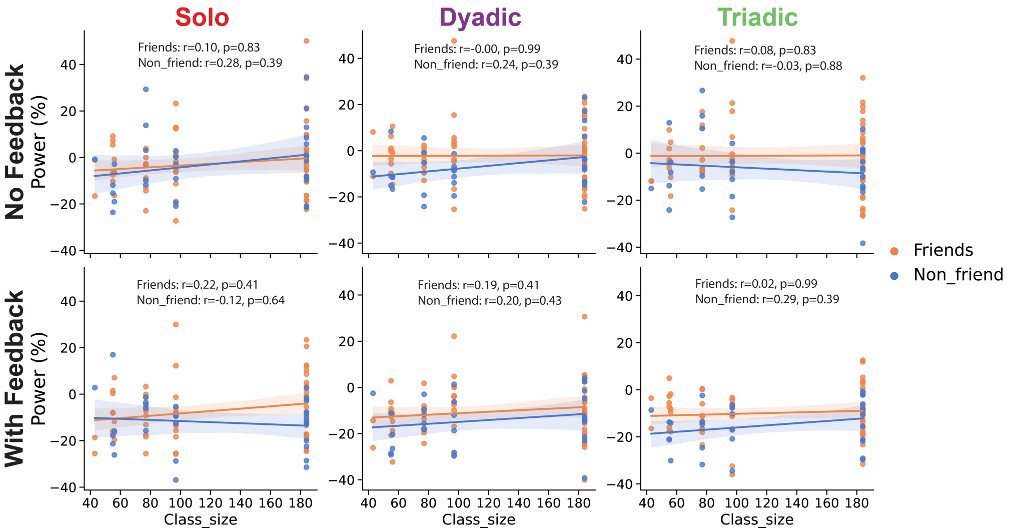


**Fig. S8**. **No correlations between beta ERD and class social network sizes.** No associations were observed between trial averaged left-sensorimotor beta ERD and the size of the class social networks. Spearman correlation coefficients were computed and the p-values shown have been FDR corrected.

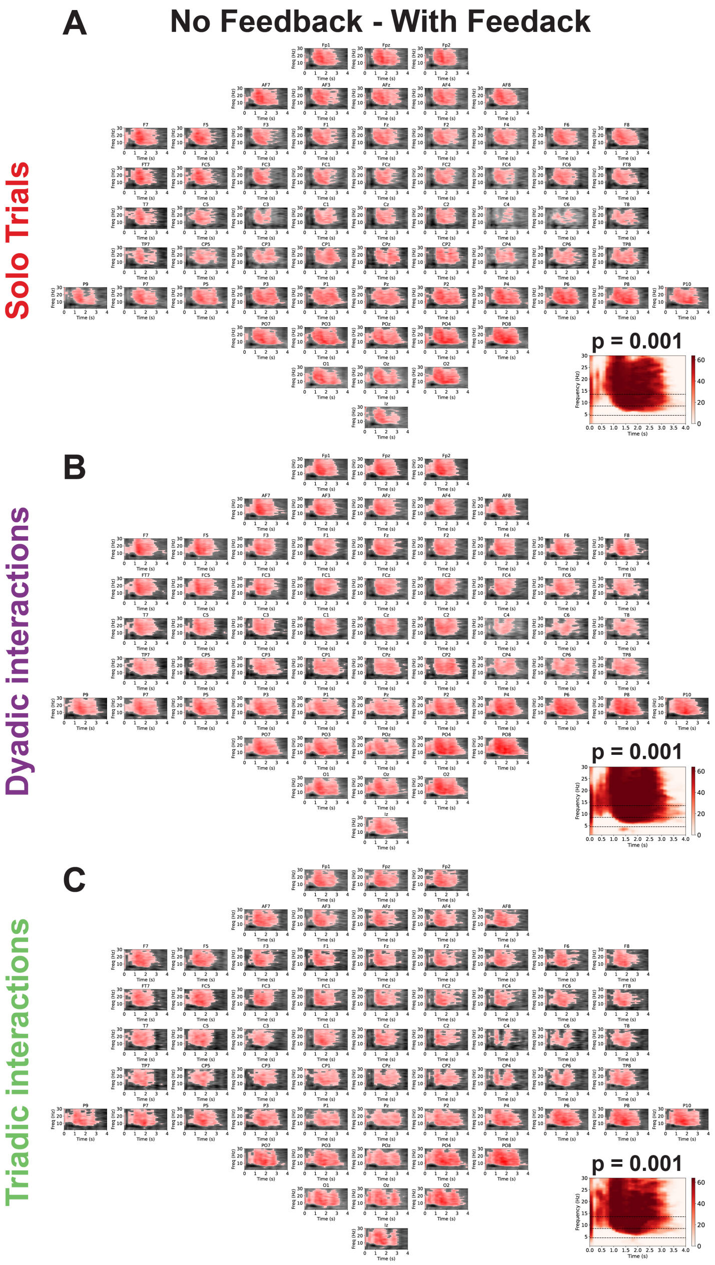


**Fig. S9**. **Non-parametric cluster-based permutation test for effect of feedback on event-related power.** T-statistics are shown in spectrograms with time on the x-axis and freq on the y-axis, with subplots for each electrode, ordered spatially with the top row corresponding to most anterior positions. Darker colors indicate greater t-values, with red colors indicating the data points are part of a significant cluster with more desynchronization in trials with continuous visual feedback. Greater desynchronization with feedback was observed in A) solo, B) dyadic and C) triadic trials. Clusters consistently found across solo, dyadic and triadic conditions spanned from approximately 0.7s to 3.0s and across theta, alpha and beta frequency ranges, globally across all electrodes. The insets show summary spectrograms collapsed over the spatial dimension (electrodes), counting in how many electrodes, each data point in the spectrograms were part of the significant cluster.

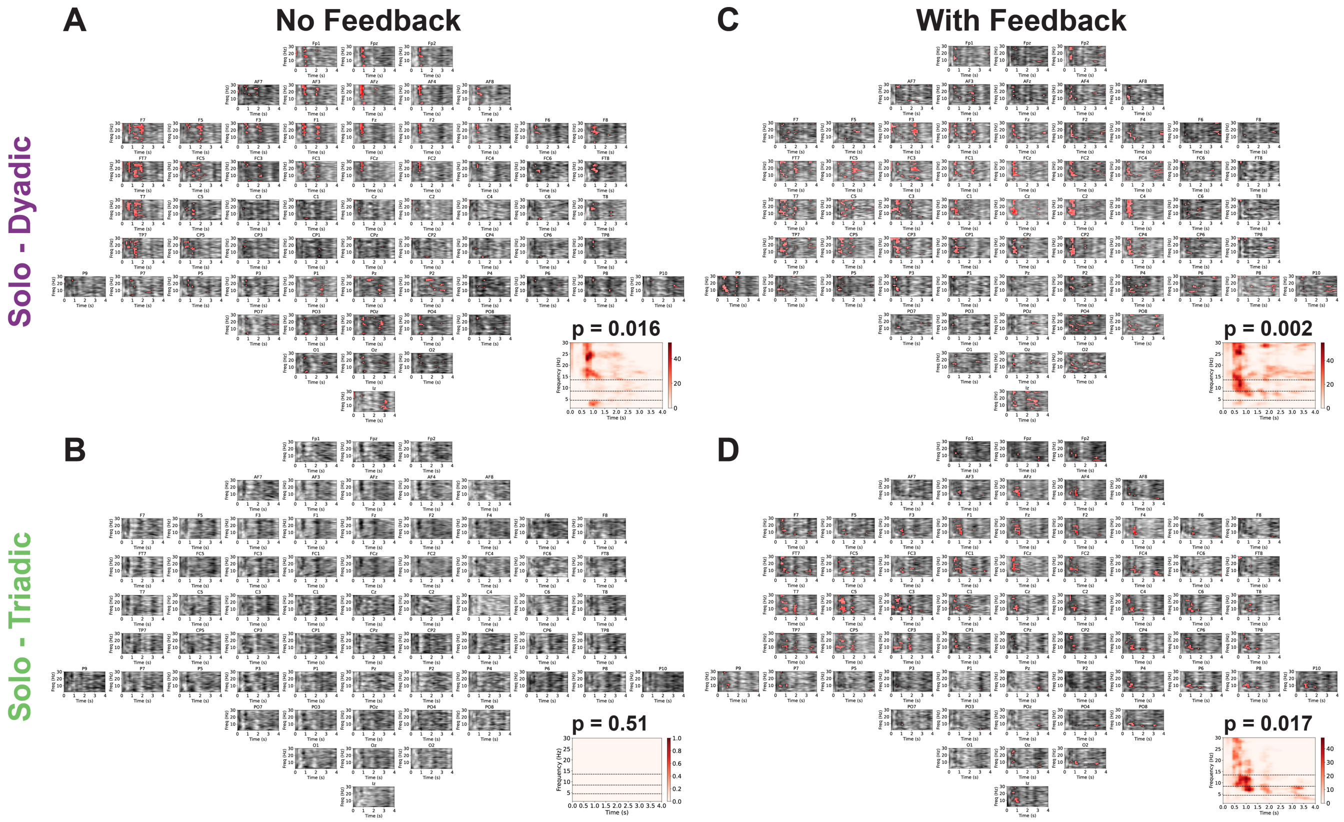


**Fig. S10**. **Non-parametric cluster-based permutation test for effect of interaction on event-related power.** T-statistics are shown in spectrograms with time on the x-axis and freq on the y-axis, with subplots for each electrode, ordered spatially with the top row corresponding to most anterior positions. Darker colors indicate greater t-values, with red colors indicating the data points are part of a significant cluster with more desynchronization in interactive group conditions compared to the solo condition. A) Dyadic trials without feedback had greater desynchronization. The significant cluster primarily covered the beta frequency band in frontal and lateral central areas, approximately from 0.8 to 1.1s. B) No difference in event-related power between solo and triadic conditions with feedback. C-D) In trials with continuous visual feedback, interactive conditions (both dyadic and triadic) had significantly greater desynchronization. The clusters spanned from approximately 0.4s to 1.0s and across alpha and beta frequency ranges, in central areas. The insets show summary spectrograms collapsed over the spatial dimension (electrodes), counting in how many electrodes, each data point in the spectrograms were part of the significant cluster.

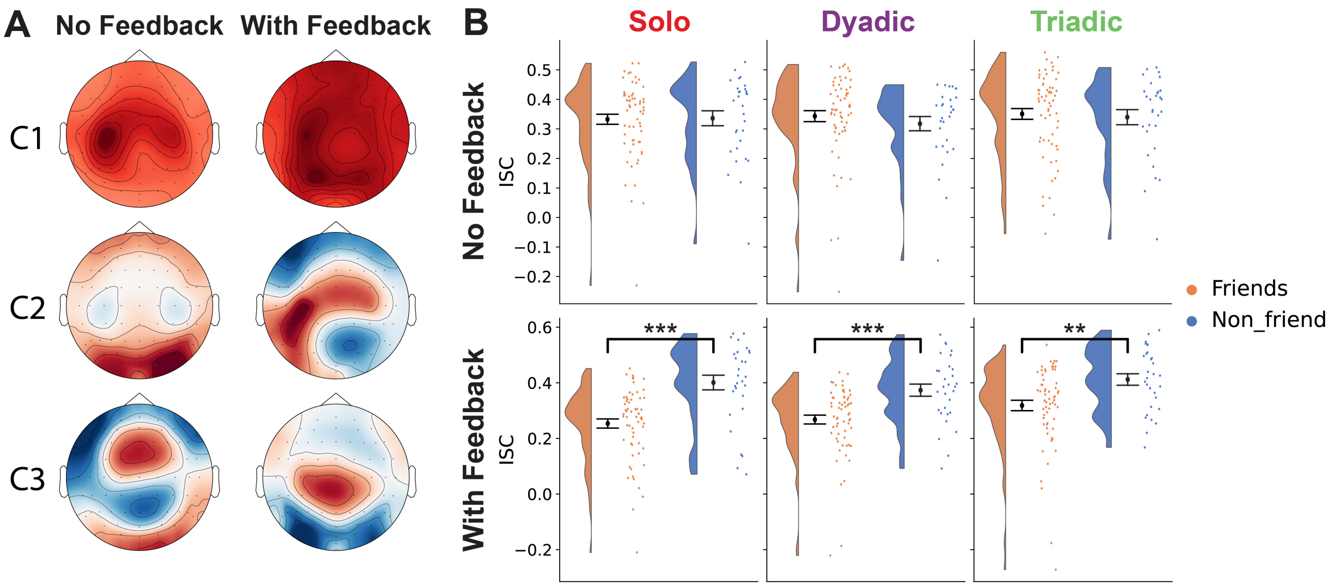


**Fig. S11**. **Non-friends exhibit increased inter-subject correlations in beta frequency range.** A) Three correlated components fitted to beta event-related power had ISC above chance levels ($p < 0.05$ estimated with circular shifted surrogate data). B) For the leading component C1, which has the highest ISC, non-friends had greater ISC compared to friends in trials with continuous visual feedback. The topographic maps correspond to projections of the forward model weights, where the polarity is normalized so the Cz electrode weight is positive. $^{**}p < 0.01$, $^{***}p < 0.001$. P-values were derived using non-parametric permutation tests with FDR multiple test correction. ISC: inter-subject correlation.

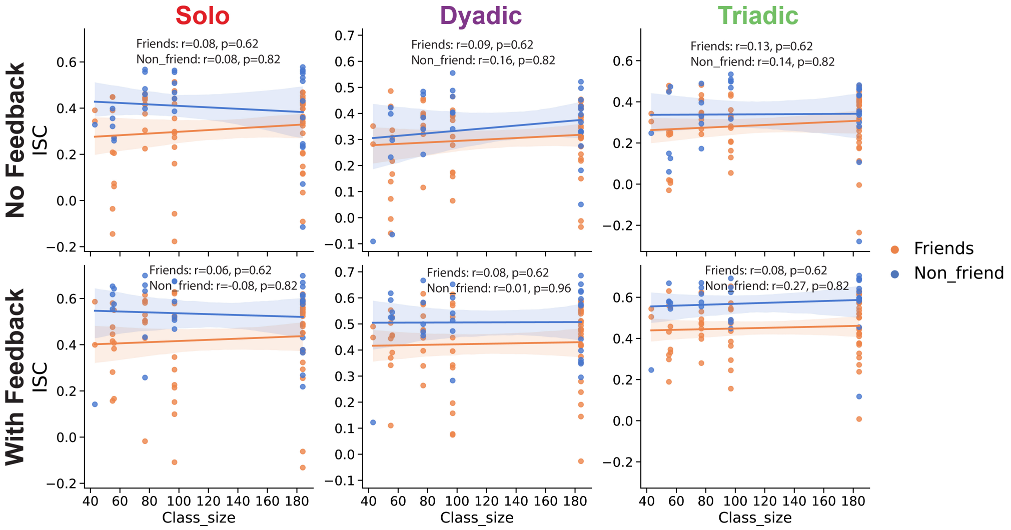


**Fig. S12**. **No correlations between alpha ISC and class social network sizes.** No associations were observed between ISC for the leading alpha component (C1) and the size of the class social networks. Spearman correlation coefficients were computed and the p-values shown have been FDR multiple test corrected.

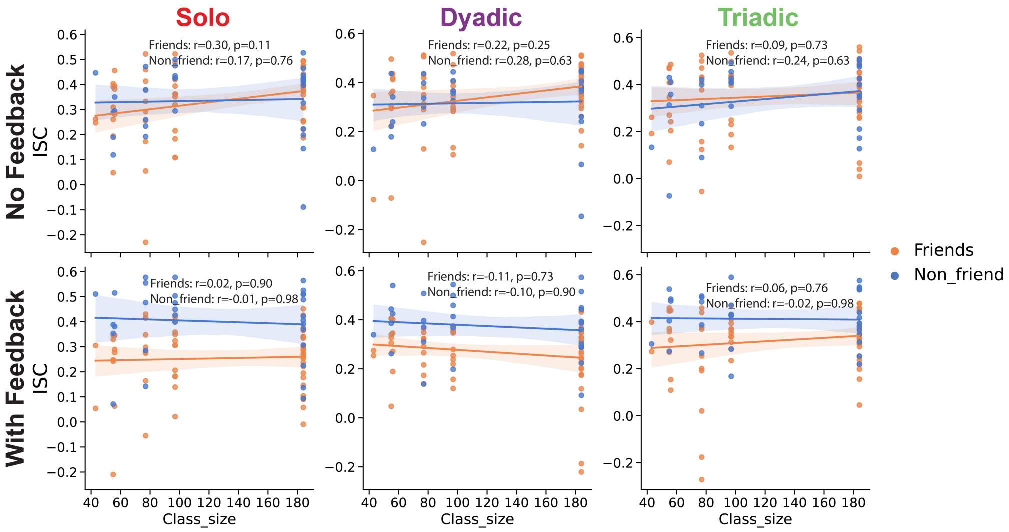


**Fig. S13**. **No correlations between beta ISC and class social network sizes.** No associations were observed between ISC for the leading beta component (C1) and the size of the class social networks. Spearman correlation coefficients were computed and the p-values shown have been FDR multiple test corrected.

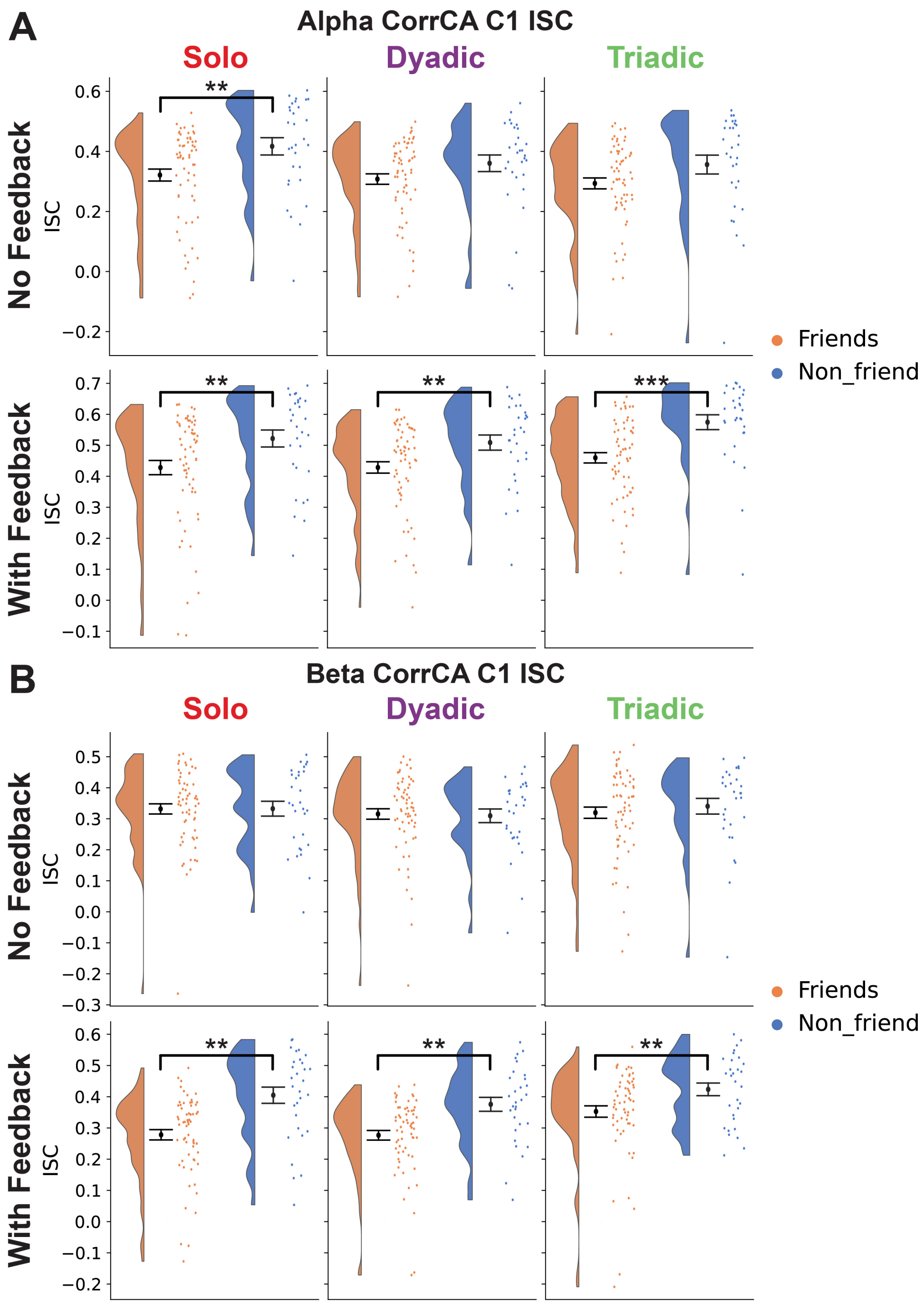


**Fig. S14**. **Non-friends still exhibit increased inter-subject correlations after standardization.** To take into account that non-friends have increased ERDs, we also tried standardizing the ERDs prior to fitting the CorrCA algorithm. The same results with increased ISC in non-friends were observed. $^{**}p < 0.01$, $^{***}p < 0.001$. P-values were derived using non-parametric permutation tests with FDR multiple test correction. ISC: inter-subject correlation.

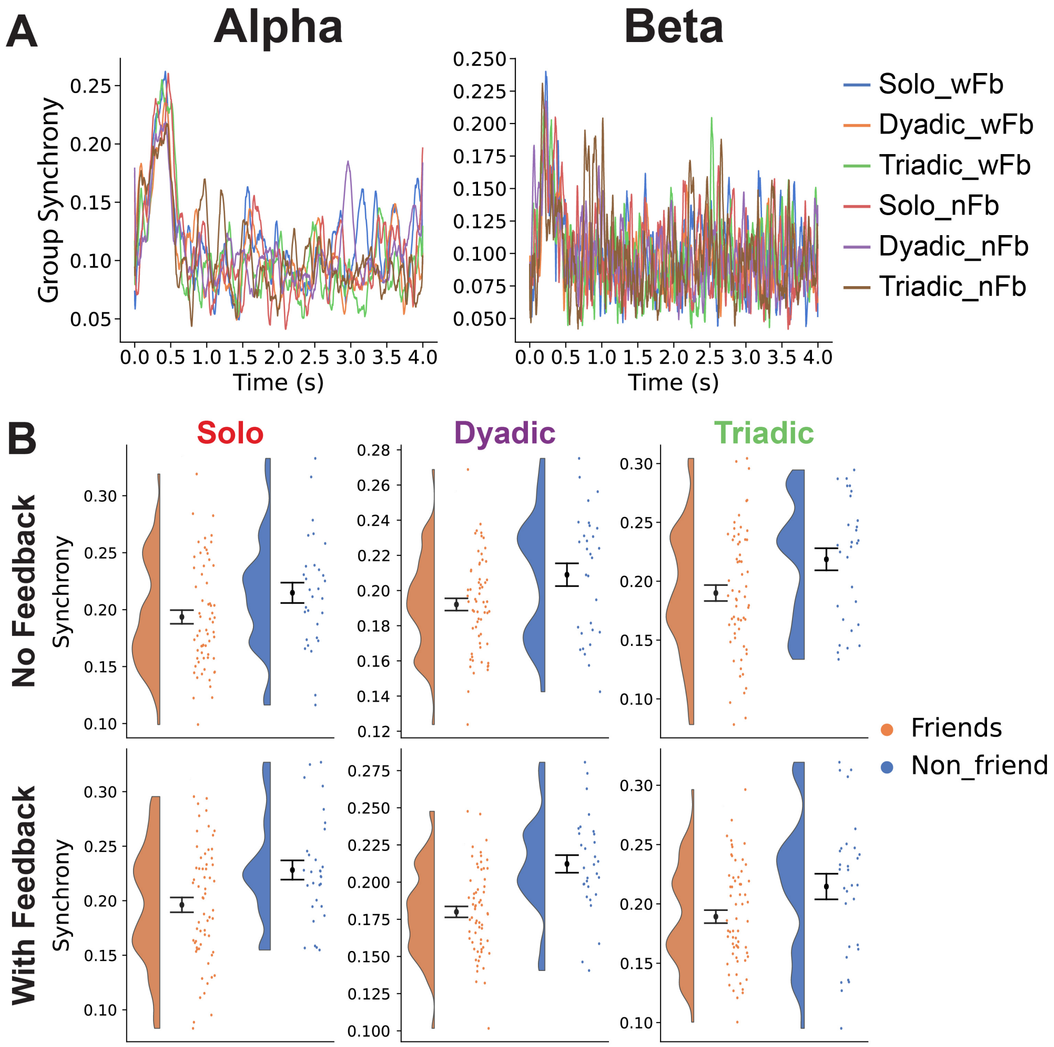


**Fig. S15**. **EEG group cluster phase synchrony.** A) Example plots of group phase synchrony from electrodes above left sensorimotor area in alpha and beta frequency bands, color-coded based on the different conditions of group sizes and feedback. B) Time averaged alpha group phase synchrony from the left sensorimotor area, plotted separately for friends and non-friends. wFB: with feedback; nFB: no feedback.

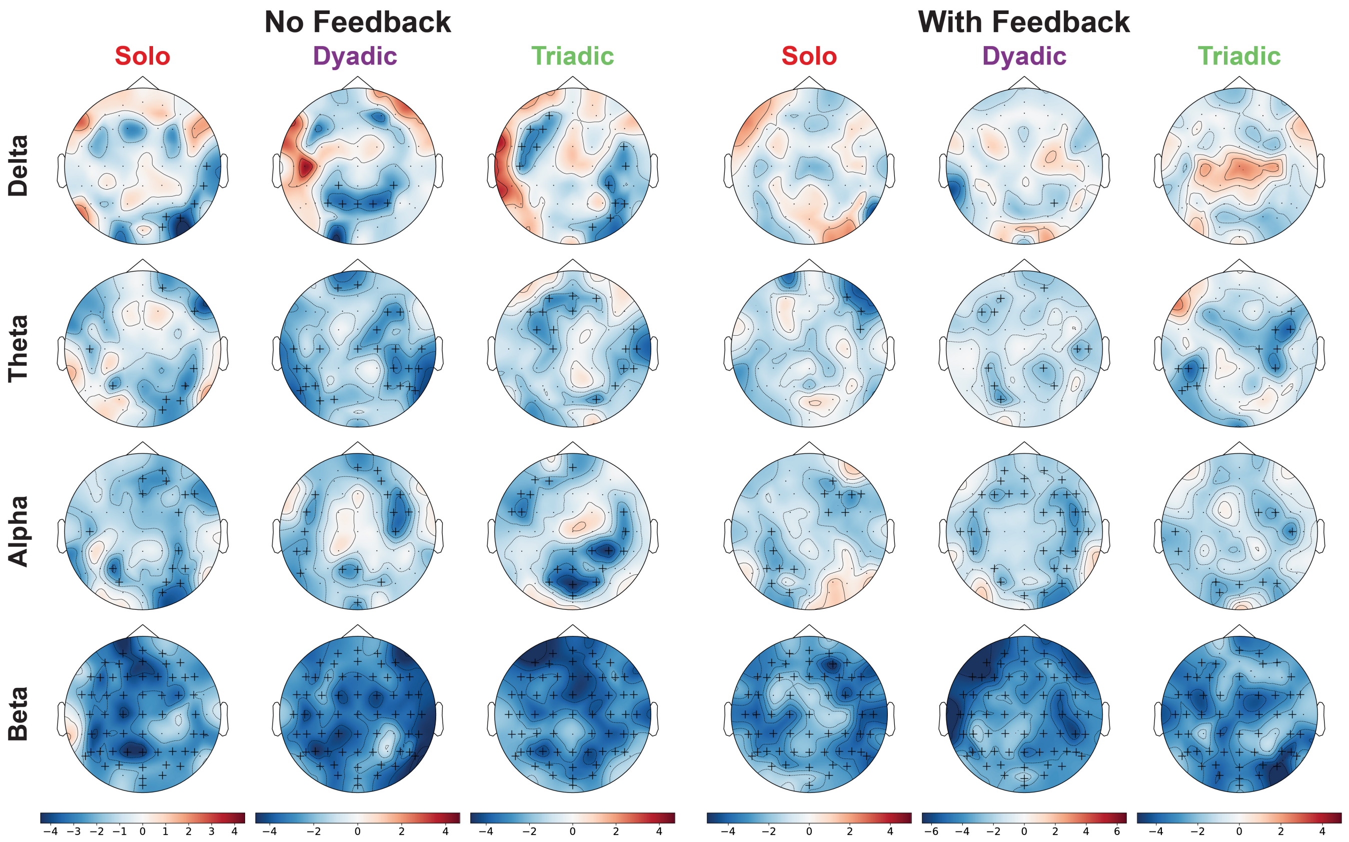


**Fig. S16**. **Non-friends exhibit increased alpha/beta group phase synchrony.** Cluster-based permutation tests of group phase synchrony across channels and frequency. Non-friends had increased group phase synchrony compared to friends in all conditions (more blue), primarily in beta and alpha frequency bands.

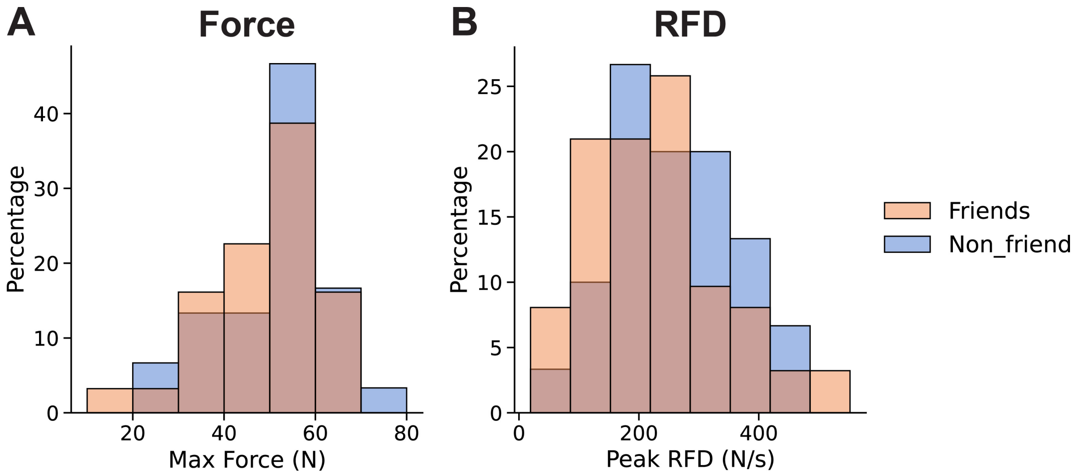


**Fig. S17**. **No effect of friendship on max force and max RFD.** All participants performed practice trials before the experiment to familiarize themselves to the devices, and in three of those trials the participants were instructed to ”Press as hard as you can on the force device after the fixation cross disappears”. These max press trials were used to get a baseline of the maximum physical strength of the participants. (A-B) Histograms showing the max force and max RFD. No difference in A) max force ($p = 0.357$) and B) max RFD ($p = 0.269$) was found between friends and non-friends. Uncorrected p-values. RFD: rate of force development.

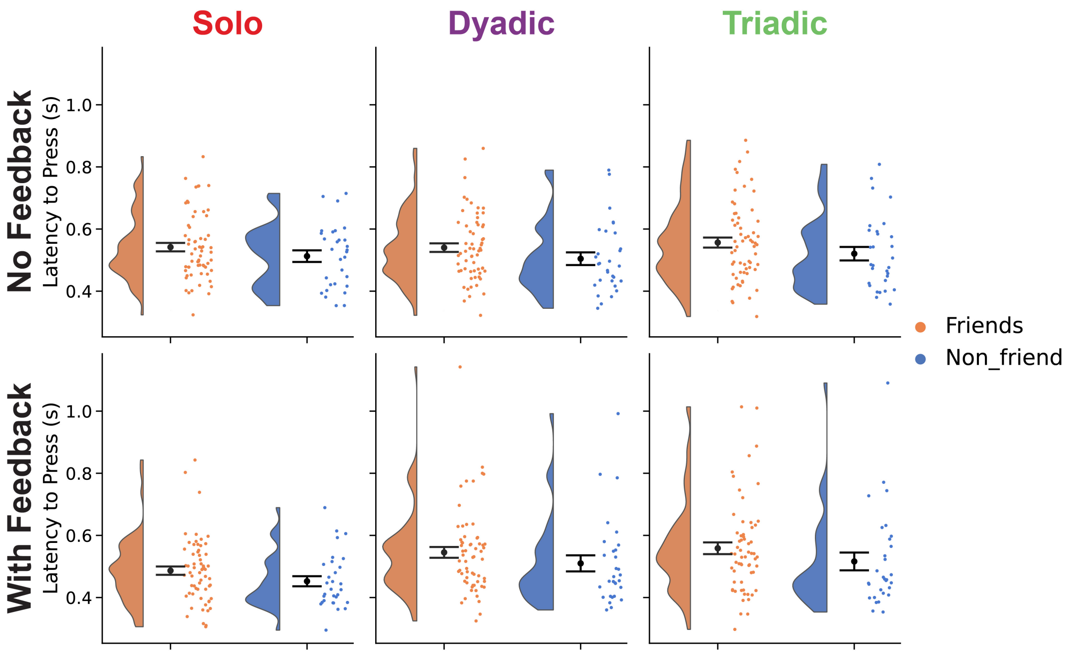


**Fig. S18**. **No effect of friendship on response times.** Response times were quantified as latency to press, after excluding false starts defined as press latencies $\leq$ 160ms, which most often corresponded to pressing before the trial started. No differences in response times were observed between friends and non-friends across all conditions ($p \geq 0.255$; FDR corrected).

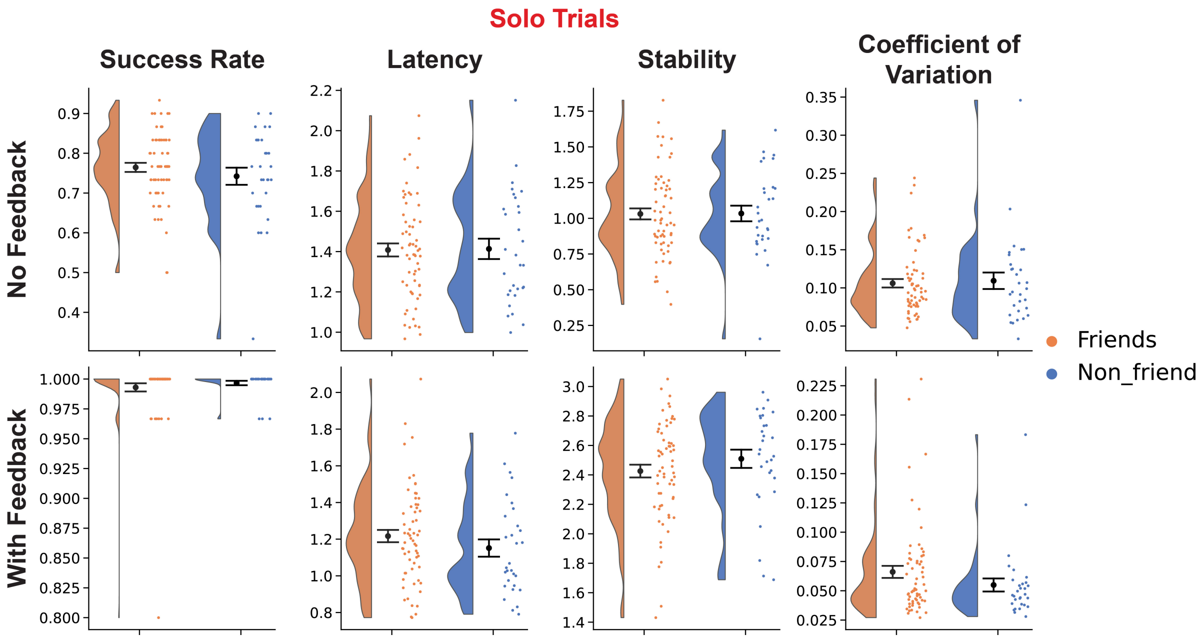


**Fig. S19**. **No effect of friendship was observed on behavioural performance in solo trials.** Success rate, latency, stability and coefficient of variation were similar for friends and non-friends during solo trials, with and without continuous visual feedback.

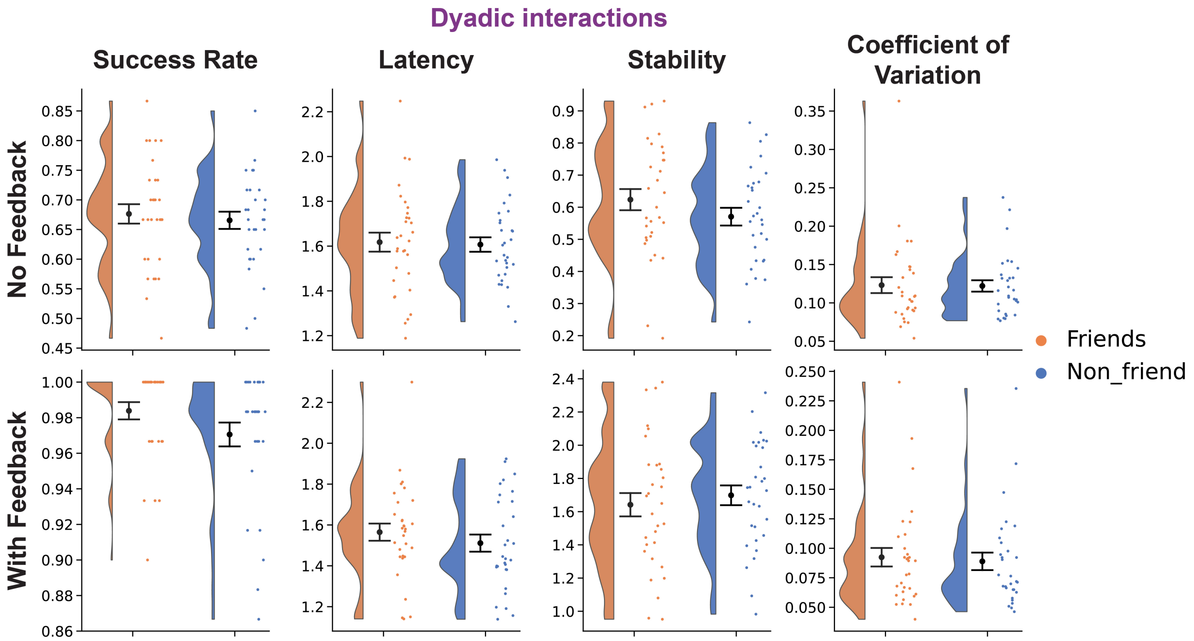


**Fig. S20**. **No effect of friendship was observed on behavioral performance in dyadic interactions.** Success rate, latency, stability and coefficient of variation were similar for dyadic trials performed by the friend pair and friend non-friend interactions, with and without continuous visual feedback.

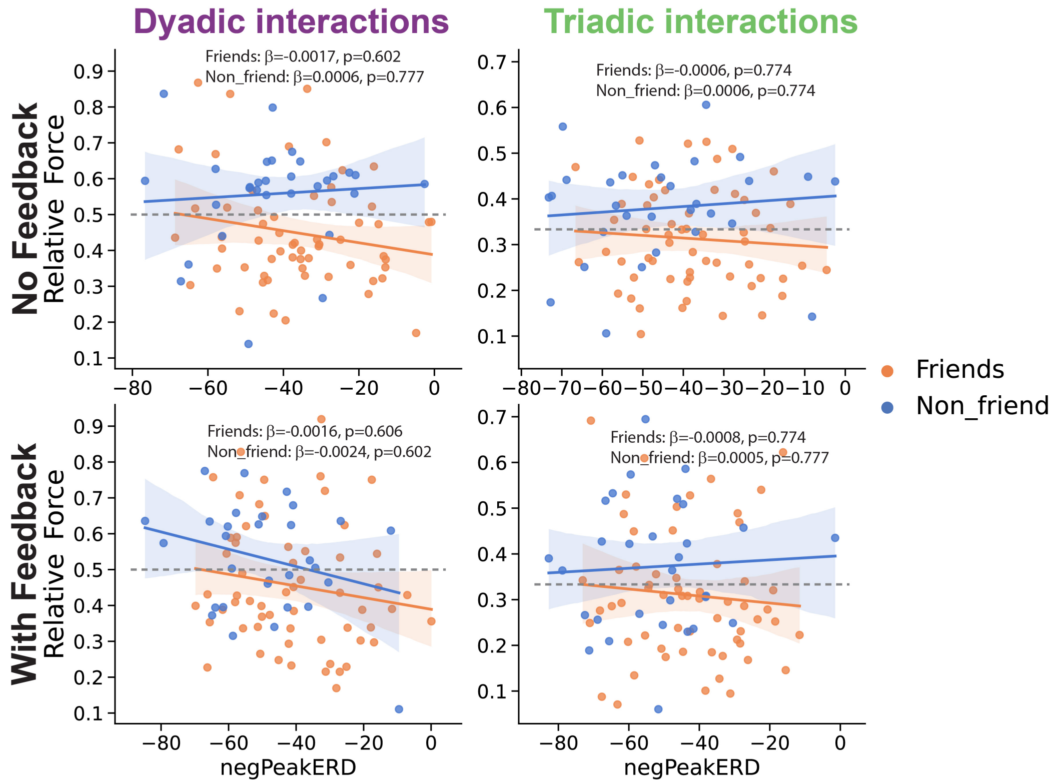


**Fig. S21**. **No association between alpha ERD magnitude and relative force.** No associations were found between the magnitude of the negative alpha ERD peak and relative force. The striped line corresponds to equal force contribution from all participants. Because the linear mixed-effects revealed no significant main effect or interaction effects involving ERD ($p > 0.12$), condition specific ordinary least square regression lines and coefficients are shown for descriptive visualization within each condition, with FDR corrected p-values. ERD: event-related desynchronization.

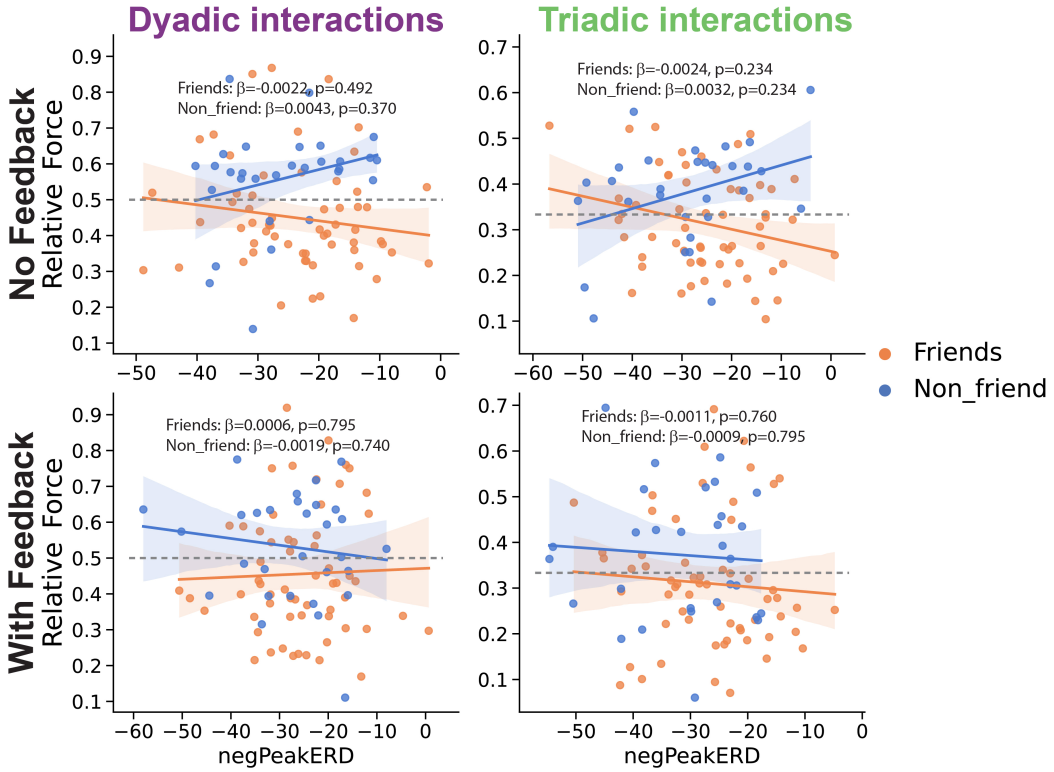


**Fig. S22**. **No association between beta ERD magnitude and relative force.** No associations were found between the magnitude of the negative beta ERD peak and relative force. The striped line corresponds to equal force contribution from all participants. Because the linear mixed-effects revealed no significant main effect or interaction effects involving ERD ($p > 0.06$), condition specific ordinary least square regression lines and coefficients are shown for descriptive visualization within each condition, with FDR corrected p-values. ERD: event-related desynchronization.

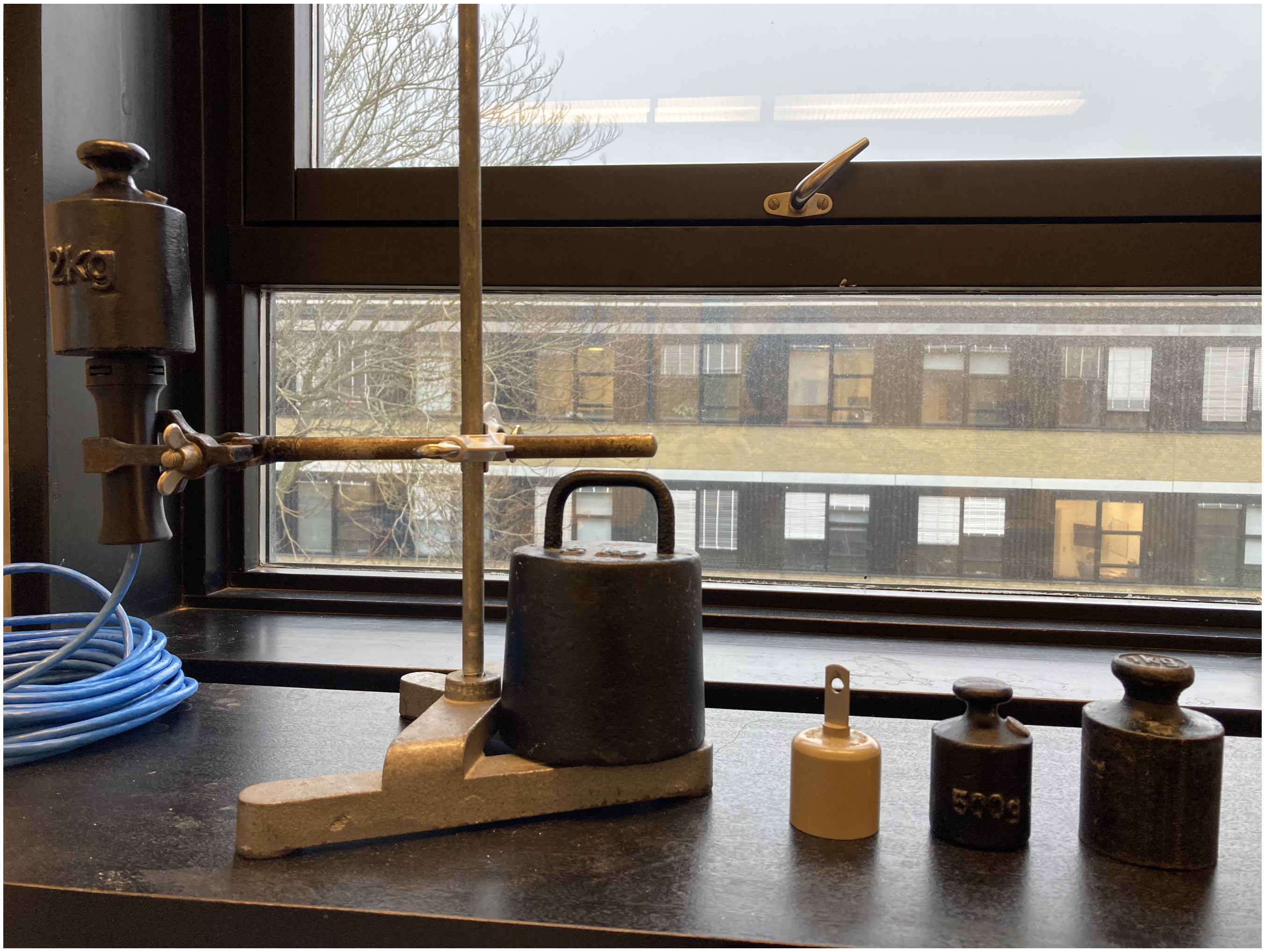


**Fig. S23**. **Calibration of the custom-made force measuring devices.** Weights with known physical weights (295g, 500g, 1000g, 2000g and 5000g) were placed stably on each force device and measured over 1 minute in order to compute the relationship between the pressure applied on each device and the digital output recorded by the computer. This enabled us to calibrate and convert the measured force into Newtons.
